# Mathematical model of intestinal lipolysis of a long-chain triglyceride

**DOI:** 10.1101/2024.05.01.592066

**Authors:** Oljora Rezhdo, Richard West, Myriel Kim, Brandon Ng, Sigal Saphier, Rebecca L. Carrier

## Abstract

Lipids are an important component of food and oral drug formulations. Upon release into gastrointestinal fluids, triglycerides, common components of foods and drug delivery systems, form emulsions and are digested into simpler amphiphilic lipids (e.g., fatty acids) that can associate with intestinal bile micelles and impact their drug solubilization capacity. Digestion of triglycerides is dynamic and dependent on lipid quantity and type, and quantities of other components in the intestinal environment (e.g., bile salts, lipases). The ability to predict lipid digestion kinetics in the intestine could enhance understanding of lipid impact on the fate of co-administered compounds (e.g., drugs, nutrients). In this study, we present a kinetic model that can predict the lipolysis of emulsions of triolein, a model long-chain triglyceride, as a function of triglyceride amount, droplet size, and quantity of pancreatic lipase in an intestinal environment containing bile micelles. The model is based on a Ping Pong Bi Bi mechanism coupled with quantitative analysis of partitioning of lipolysis products in colloids, including bile micelles, in solution. The agreement of lipolysis model predictions with experimental data suggests that the mechanism and proposed assumptions adequately represent triglyceride digestion in a simulated intestinal environment. In addition, we demonstrate the value of such a model over simpler, semi-mechanistic models reported in the literature. This lipolysis framework can serve as a basis for modeling digestion kinetics of different classes of triglycerides and other complex lipids as relevant in food and drug delivery systems.

## 1 Introduction

Lipids can significantly impact the intestinal solubility and overall absorption of orally delivered compounds, either as formulation components in delivery systems or when co-administered within lipid-rich foods (Rezhdo et al., 2016). Lipid use in oral delivery of compounds is limited in part due to lack of mechanistic understanding of and ability to predict lipid function in the gastrointestinal (GI) tract. For example, the effect of lipid type and lipid chain length on quantitative features (e.g., effective volume) of colloidal structures formed during digestion, and the impact of these structures on compound solubility and absorption across the intestinal barrier, are poorly understood (Dahan & Hoffman, 2007; Grove et al., 2006; Myers & Stella, 1992; Porter, Kaukonen, Boyd, et al., 2004; Porter, Kaukonen, Taillardat-Bertschinger, et al., 2004). Ingestion of lipids triggers a cascade of processes, including lipolysis, that change the physical and chemical nature of the gastrointestinal milieu and directly affect the fate of oral compounds in the GI tract. The majority of lipolysis occurs in the small intestine (Amara et al., 2019) primarily because oil droplets are emulsified by bile salts upon entering the duodenum, resulting in an increase in the available surface area over which lipolysis can occur (Maldonado-Valderrama et al., 2011). The mechanism by which lipids are digested by lipases is complex and not fully understood (Aloulou et al., 2008), in part due to lipolysis occurring at the oil-water interface vs. in solution and the associated challenge of quantifying lipid and lipase content at the interface. Various kinetic models have been proposed (Al-Zuhair et al., 2003; Al-Zuhair et al., 2004a, 2004b; Fadiloğlu & Söylemez, 1997; Garcia et al., 1992; Goto et al., 1992; Kawano, Kawasaki, et al., 1994; Kawano, Kiyoyama, et al., 1994; Martinez et al., 1992; Mohapatra & Hsu, 1997; Paiva et al., 2000; Plou et al., 1996; Rice et al., 1999; Shiomori et al., 1995; Tanigaki et al., 1995; Tsai et al., 1991; Verger et al., 1973; Wang et al., 1985). Some of these models express the rate of hydrolysis at the oil-water interface as a simple function of the interfacial area available for digestion (Buyukozturk et al., 2013; Giang et al., 2016; Giang et al., 2015; Li & McClements, 2010). This approach offers the advantage of relatively simple equations with few kinetic coefficients, and model fitting and estimation of kinetic coefficients are consequently relatively straightforward. However, these simple models do not account for concentration of enzyme in solution, limiting the ability to predict lipolysis kinetics *in vivo* where enzyme concentration changes during intestinal transit. These simple models also do not account for the presence of intermediates, such as diglycerides (DG) and monoglycerides (MG) produced from triglyceride (TG) digestion. Ingested triglycerides are partially hydrolyzed by gastric lipases into DG and fatty acid (FA), and thus a mixture of TG, DG, and FA transits into the small intestine (Bakala N’Goma et al., 2012; Carriere et al., 1993). The ability to capture the hydrolysis of DG (as well as MG) in addition to TG during intestinal lipolysis as well as enzyme concentration would allow for more complete prediction of the composition of the intestinal milieu, and thus understanding of drug solubilization capacity over time after TG ingestion. Models incorporating more mechanistic aspects of lipolysis at interfaces are inherently more complex and include a significant number of kinetic coefficients, thus requiring extensive data sets for model training and testing. Multiple models have been proposed for hydrolysis of lipids and used to model data from experiments incorporating different oil-water interface configurations (e.g., different reactor designs). The Ping Pong Bi Bi mechanism, the most accurate and widely accepted description of catalytic action of lipases as reviewed by Paiva et al. (Paiva et al., 2000), has been used to represent interfacial TG lipolysis including formation of intermediate species (Hermansyah et al., 2006; Hermansyah et al., 2010; Mitchell et al., 2008; Pilarek & Szewczyk, 2007). A Ping Pong Bi Bi mechanism as applied to TG describes a double displacement reaction that requires two substrates and yields two products (Figure 1). The general Ping Pong Bi Bi-based model as described by Paiva et al. is complex and contains many coefficients; thus, a roadmap for potential experimental conditions and assumptions that lump various parameters into a few groups is presented. Others have adapted Ping Pong Bi Bi-based models to their specific systems. For example, Pilarek et al. developed a model for the alcoholysis of TG in propanol by 1,3-specific lipase immobilized on a resin, including MG isomerization and irreversible enzyme deactivation (Pilarek & Szewczyk, 2007). Mitchell et al. developed a model for stepwise alcoholysis of triolein by lipase assuming an enzyme specificity constant for each substrate (i.e., TG, DG, MG) that does not change with changing physicochemical properties of emulsions during digestion (Mitchell et al., 2008). Hermansyah et al. developed a model for the stepwise hydrolysis of triolein by lipase considering partitioning of digestion products between a planar reaction interface with constant interfacial area and an oil phase (Hermansyah et al., 2006). This kinetic model was applied to experimental profiles of lipid species measured from the oil phase of the solution.

**Figure 1:**
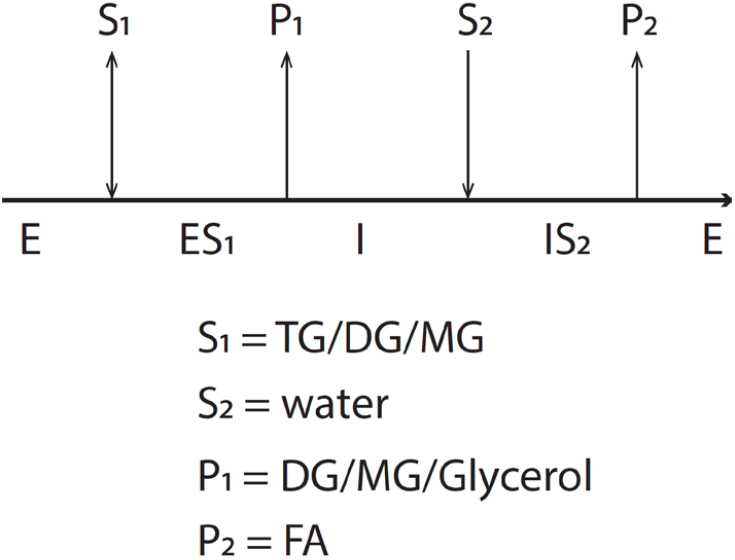
Schematic representation of a general Ping-Pong Bi Bi mechanism in the context of one of the steps of triglyceride hydrolysis. S_1_ and S_2_ represent substrates, E is the enzyme, P_1_ and P_2_ are products. ES_1_ and IS_2_ represent enzyme-substrate complexes. I represents an enzyme-fatty acid intermediate which requires a water molecule before FA can be released.

In this study, we built upon the original model proposed by Hermansyah et al. (Hermansyah et al., 2006) and adapted it to incorporate essential features of the intestinal environment to enable prediction of lipolysis kinetics in a physiological context. Specifically, as the mixed intestinal environment after lipid ingestion consists of emulsions evolving in size due to lipolysis, we accounted for changing spherical interfacial area as a function of the quantities of lipid species during lipolysis. Additionally, we adapted the model to capture the total concentration of lipid species at the oil droplet interface vs. the droplet core or the solution surrounding the droplets by considering a portion of the digestion products dissociating from the interface (e.g., by partitioning into the oil core, by associating with micelles and/or forming vesicles in the aqueous phase). It is noted that knowledge of the concentrations of lipid digestion products associating with colloidal species in the bulk solution is important as it impacts the solubilization capacity of the intestinal milieu, and associated drug dissolution kinetics and solubility. Experiments conducted to test the model included dispersion of triglyceride to form emulsion droplets in simulated intestinal fluids containing model bile components, such that the lipid phase consisted entirely of TG and digestion products. Further, the pH of the lipolysis reaction was maintained via titration. Thus, the model was tested using an experimental system that captured essential features of the native intestinal environment absent from previous studies, which did not contain model bile and allowed lipolysis to occur at a planar interface between an organic solvent phase and an aqueous phase (Hermansyah et al., 2006; Hermansyah et al., 2010). Triolein (TO, C18:1) was studied as a representative long-chain triglyceride. We subjected the model to training and test sets of kinetic data to determine model coefficients and validate the predictive ability of the model for triolein emulsion digestion, respectively, in the simulated mixed intestinal environment including bile micelles. Lastly, we compared the proposed model to other intestinal lipolysis models discussed above to explore its benefits and drawbacks.

## 2 Materials and Methods

### 2.1 Materials

Trizma maleate (P/N: T3128), sodium chloride (NaCl, P/N: 71376), calcium chloride dihydrate (CaCl_2_ · 2H_2_O, P/N: C5080), sodium azide (NaN_3_, P/N: S2002), sodium taurodeoxycholate hydrate (NaTDC, P/N: T0557), L-alpha-phosphatidylcholine from egg yolk (EYPC, P/N: P3556), glyceryl trioleate or triolein (TO, P/N: T7140), 1-oleoyl-rac-glycerol (MO, P/N: M7765), oleic acid (OA, P/N: O1008), and pancreatin from porcine pancreas 8 x USP (P/N: 7545) were purchased from Sigma Aldrich Inc., St. Louis, MO, USA. Diolein (DO, P/N: D-236-N9-Z) was purchased from Nu-Chek Prep Inc., Waterville, MN, USA. Coconut oil (P/N: CO110) and palm oil (P/N: PA105) were purchased from Spectrum Chemical, New Brunswick, NJ, USA. Linseed oil was purchased from MP Biomedicals, Santa Ana, CA, USA (P/N: 960122). Virgin olive oil was purchased from Thermo Scientific, Waltham, MA, USA (P/N: 41654). Acetone (P/N: A18P), acetonitrile (P/N: A998), methanol (P/N: A998), and chloroform (P/N: C607) were purchased from Thermo Fisher Scientific Inc., Waltham, MA, USA. The HPLC 1260 in line with a charged aerosol detector (CAD) was obtained from Agilent Technologies, Santa Clara, CA, USA. The CAD detector was obtained from formerly ESI Biosciences, now part of Thermo Fisher Scientific Inc., Waltham, MA, USA. The TL-100 ultracentrifuge was obtained from Beckman Coulter, Brea, CA, USA. The Litesizer 100 particle size analyzer was obtained from Anton Paar, Graz, Austria. An Omni Sonic Ruptor 400 Ultrasonic Homogenizer, 115V was obtained from Omni International The Homogenizer Company, Kennesaw, GA, USA. Computational modeling and data fitting were performed using MATLAB R2016A (9.0.0.341360), MathWorks, Natick, MA, USA.

### 2.2 *In vitro* simulation of intestinal digestion

The composition of simulated intestinal fluids containing model bile micelles at fed concentrations is summarized in Table 1 (Di Maio & Carrier, 2011). The fed state solution was prepared for all studies reported in this article at a 4X concentration to allow room for dilution to a final 1X concentration upon addition of enzymatic cocktail and test lipid. The solution was prepared fresh the day before the experiment and mixed overnight at 37°C. The bile concentrations selected here are representative of values determined in the human duodenum in the fasted state and after a standard hospital diet (Mallory et al., 1973), and the simulated intestinal fluid compositions employed are broadly used in pharmaceutical research, including lipid digestion studies, for representing intestinal contents (Minekus et al., 2014; Williams, Anby, et al., 2012; Williams, Sassene, et al., 2012; Williams et al., 2013).

**Table 1:** Composition of simulated intestinal fluids (SIF) containing bile micelles at fed concentrations (FeSSIF) and fasted concentrations (FaSSIF)

|  | **Compound** | **Concentration** |
| --- | --- | --- |
| **Simulated Intestinal Fluids (SIF)** | Trizma Maleate | 100mM |
|  | NaCl | 65mM |
|  | CaCl<sub>2</sub> · 2H<sub>2</sub>O | 10mM |
|  | NaN<sub>3</sub> | 3mM |
| **Model bile micelles at <u>fed</u> concentrations** | NaTDC | 12mM |
|  | EYPC | 4mM |
| **Model bile micelles at <u>fasted</u> concentrations** | NaTDC | 3mM |
|  | EYPC | 1mM |

Pancreatin extract was prepared by adding 2 g pancreatin to 10 mL simulated intestinal fluids and mixing at room temperature for 15 min. The solution was then ultracentrifuged at 76,000 rpm for 30 min. The supernatant containing the pancreatin extract was collected and stored at 4°C until use. The extract was used within 5 h from preparation. The activity of the extract was measured as recommended by Carriere et al. (Carriere et al., 1993). Lastly, a physiological range of concentrations for TO was selected based on values from human intestinal content measurements by Armand et al. (Armand et al., 1994). The conditions for *in vitro* intestinal digestion are shown in Table 2.

**Table 2:** List of experimental conditions used for simulation of intestinal digestion in vitro. Experiments 1 – 10 are conducted using FeSSIF media. Experiments 11 – 12 are conducted using FaSSIF media. TO = triolein. Concentrations of TO shown are in final solution (lipid-rich solution + pancreatin extract)

| **Purpose** | **No.** | **Lipid Concentration (in 20 mL final volume)** | **Pancreatin extract\*\*** |
| --- | --- | --- | --- |
| **Training set** | 1 | 25 mM TO + FeSSIF 4X (5 mL) + water (9 mL) | 6 mL |
|  | 2* | 25 mM TO + FeSSIF 4X (5 mL) + water (9 mL) | 6 mL |
|  | 3* | 25 mM TO + FeSSIF 4X (5 mL) + water (9 mL) | 6 mL |
|  | 4 | 25 mM TO + FeSSIF 4X (5 mL) + water (13 mL) | 2 mL |
|  | 5 | 40 mM TO + FeSSIF 4X (5 mL) + water (13 mL) | 2 mL |
|  | 6 | 70 mM TO + FeSSIF 4X (5 mL) + water (9 mL) | 6 mL |
|  | 7* | 70 mM TO + FeSSIF 4X (5 mL) + water (9 mL) | 6 mL |
| **Test set** | 8 | 10 mM TO + FeSSIF 4X (5 mL) + water (13 mL) | 2 mL |
|  | 9 | 20 mM TO + FeSSIF 4X (5 mL) + water (9 mL) | 6 mL |
|  | 10 | 50 mM TO + FeSSIF 4X (5 mL) + water (9 mL) | 6 mL |
| **Training set** | 11 | 25 mM TO + FaSSIF 4X (5 mL) + water (13 mL) | 2 mL |
| **Test set** | 12 | 5 mM TO + FaSSIF 4X (5 mL) + water (13 mL) | 2 mL |

\*Same conditions as previous experiment but with larger emulsion sizes (i.e., homogenized for less time)
\*\*6 mL pancreatin extract in 20 mL total volume corresponds to a lipase activity of 1800 U/mL whereas 2 mL pancreatin extract in 20 mL final volume corresponds to a lipase activity of 600 U/mL

Test “meals” containing triolein were mixed with FeSSIF or FaSSIF media, depending on the experiment, and water, and were homogenized using an ultrasonic homogenizer at varying pulse strength and duration, depending on the lipid concentration to achieve an oil emulsion size less than 10 μm, which is the maximum size limit detectable by the Litesizer 100. Intestinal digestion was started by mixing lipid solution with pancreatin extract, generating a final pancreatin activity of 1800 U/mL solution volume (when mixing 6 mL pancreatin into 20 mL final volume) or 600 U/mL (when mixing 2 mL pancreatin into 20 mL final volume), where 1 U corresponds to 1 μmol of free FA released per minute from a substrate of tributyrin at 37°C and pH 8.0 (Minekus et al., 2014). These pancreatin activities are in the range of those measured in human intestinal contents after administration of a 960 kcal meal (Armand et al., 1994). The pH was adjusted manually to maintain pH 6.5 using 0.2M NaOH over a period of 3h. Samples were collected over time to analyze the lipid content during digestion. A total of 46 time points were collected in experiments used for model training and a total of 12 time points were collected in experiments used for model testing. To quantify the lipid profile, 100 μL of sample from the digesting solution was mixed with 900 μL of a 1:1 methanol/chloroform solution. This step ensured formation of a single phase, precipitation of lipase out of solution and prevention of further lipolysis. The samples were centrifuged at 16,000 x g at 4°C for 15 min and the supernatant was transferred into HPLC vials for lipid quantification. The HPLC conditions used are described in section 2.3.

### 2.3 Quantification of lipid profile during digestion

Quantification of lipid concentration in solution was performed via HPLC in line with a CAD detector based on a modification of the method of Olsson et al. (Olsson et al., 2014). Specifically, samples were run through a ProntoSIL 5 μm column (250 x 46mm) using a 1 mL/min 5mM ammonium acetate in water-5mM ammonium acetate in methanol gradient method as shown in Table 3. Samples were prepared in 1:1 methanol/chloroform solution and 5 μL sample was injected. A quadratic least squares regression model was fit to each of the lipid standard curves. All standard curves generated a correlation coefficient R^2^ > 0.99.

**Table 3:** HPLC conditions for quantifying lipid profile

|  |  |
| --- | --- |
| Solvent A: | 5mM Ammonium Acetate + water |
| Solvent B: | 5mM Ammonium Acetate + methanol |
| Detection method: | CAD |
| Timetable: | 0 min: 30% A – 70% B<br>2 min: 30% A – 70% B<br>20 min: 0% A – 100% B<br>30 min: 0% A – 100% B<br>31 min: 30% A – 70% B<br>40 min: 30% A – 70% B |

### 2.4 Quantification of oil emulsion/particle size

Triolein emulsion droplet size was analyzed using an Anton Paar Litesizer 100 at 37°C. The solution refractive index was set to 1.340, viscosity was set to 0.691 cP, and the scattering angle was at 175.0°, allowing measurement of undiluted samples to avoid dilution artifacts. Samples collected from the digesting solution or prior to start of digestion were placed in disposable cuvettes and analyzed. Measurements consisted of 1 min temperature equilibration followed by approximately one minute optics adjustment. A maximum of 60 measurements at 10s per measurement were conducted until a scattering intensity was achieved that was adequate for particle distribution analysis. On average, one full sample measurement took 5 min. Analysis of pancreatin extract (at 3.3-fold more concentrated than during digestion) and the FeSSIF media (at 4-fold more concentrated than during digestion) indicated presence of particles in the 5-15 nm range, which likely correspond to micelles. Despite these solutions being significantly more concentrated than during lipolysis, their light transmittance recorded by the instrument was about 70%, contrary to the oil emulsions where the transmittance was about 0.1%. These observations suggest that micellar particles are dilute relative to the oil particles in a homogenized triolein in FeSSIF solution, and that the presence of micelles in solution does not affect the analysis of oil emulsion size.

### 2.5 Lipolysis kinetic measurements for different triglycerides

To evaluate the similarity of digestion kinetics of triolein, a pure triglyceride containing a single type of fatty acid, relative to naturally occurring long-chain triglycerides we conducted digestion experiments using 20mM triglyceride (concentration in final volume) + 5mL FeSSIF (4X) + 9mL water + 6 mL pancreatin. Five triglycerides were tested, including triolein, virgin olive oil, linseed oil, palm oil, and coconut oil. The molecular weight of each triglyceride was calculated based on the fatty acid composition provided either by the vendor or from the literature. A weighted average fatty acid molecular weight was computed which was converted to a triglyceride molecular weight using the formula 3 x *MW*_*FA*_ + *MW*_*Glycerol*_ – 3 x *MW*_*water*_. To evaluate the digestion kinetics of the complex lipids (i.e., consisting of a mixture of different types of fatty acids), the solutions were titrated to pH 6.5 using a pH stat autotitrator and 0.2M NaOH. The amount of NaOH added was used to determine the amount of FA released. The experiments were allowed to continue for 120 min and NaOH was added as needed. Triolein digestion was analyzed using the titration method as well as by HPLC (section 2.3) for comparison of the two analytical techniques.

## 3 Proposed mathematical model of lipolysis in the intestine

Hydrolysis of TG occurs at the oil-water interface where pancreatic lipases adsorb and attack lipids to produce DG, MG, and FA. Here, it is assumed that no long chain lipid species are free in water, given that their aqueous solubilities are negligible relative to other concentrations in solution (Labourdenne et al., 1996). Long chain TG and DG are known to not associate with micelles in solution (Reis et al., 2009; Reis et al., 2008). They are thus considered to be only at the oil-water interface and at the core of the oil droplets (Figure 2). As triolein is a liquid long-chain triglyceride, diffusion from the interface into the core and thus partitioning of TO and DO between these two phases is considered instantaneous relative to other digestion-related processes, as assumed in other lipolysis models in the literature (Hermansyah et al., 2006). In contrast, MO and OA formed at the oil-water interface by hydrolysis of TO and DO, respectively, are considered swelling amphiphiles and are known to partition between the lipid-aqueous interface and the micelles in the aqueous phase (Rezhdo et al., 2017). Partitioning of these two amphiphiles into the oil core is assumed negligible due to the hydrophilic glycerol backbone of MO and carboxyl group of OA (Figure 2). Monoglycerides are known to significantly decrease the oil interfacial tension and expel other adsorbed amphiphilic components from the interface (Reis et al., 2008), and are assumed to instantaneously partition between the interface and the aqueous phase. OA is expected to follow a similar partitioning mechanism. Considering the large amount of OA formed during digestion of TO (Figure 4), it was of interest to understand what fraction of OA could partition into the interface. A study by Couallier et al. showed that the area occupied by one OA molecule at an oil-water interface was approximately 36 Å^2^ (Couallier et al., 2018). The amount of OA needed to cover the entire oil surface area for a 50mM triolein solution consisting of 1 μm diameter particles would thus be 1.3mM. Upon initiation of hydrolysis, the amount of TO and thus the capacity of its interface to “carry” OA depletes. Additionally, the interface contains a mixture of DO and MO molecules, thus further decreasing its capacity for OA. From our data we observe a high concentration (>10 mM) of OA measured in solution within minutes of initiation of digestion (Figure 4). Thus, it is likely that most, if not all, of the OA formed partitions away from the interface (Figure 2).

**Figure 2:**
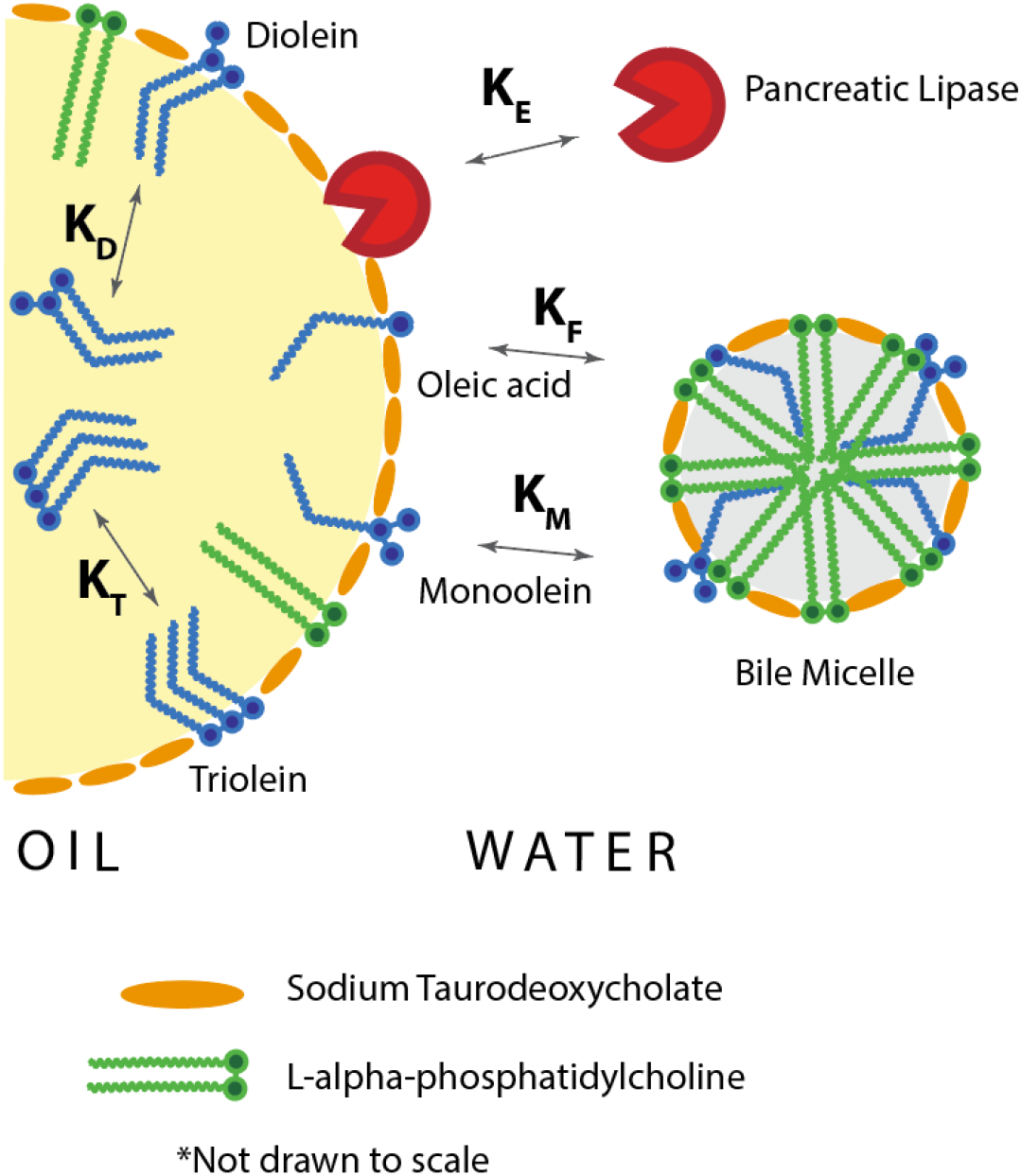
Proposed distribution of lipolysis species in the oil-water mixture where TO and DO are found only in the oil core and oil-water interface whereas MO, OA, and enzyme are found in the micelle-rich water phase and at the oil-water interface. K, K_D_, K_M_, K_F_, and KE represent the partition coefficients of TO, DO, MO, OA, and enzyme, respectively.

Under these assumptions, we propose an interfacial stepwise kinetic model following the Ping Pong Bi Bi mechanism for TO, DO, and MO hydrolysis as shown in Figure 3. As summarized in Table 4, T, D, M, F, G, E, and W correspond to triolein, diolein, monoolein, oleic acid, glycerol, enzyme, and water, respectively. The asterisk refers to the compound at the oil-water interface. E*T*, E*D*, E*M*, and E*F* correspond to lipid-enzyme complexes reversibly formed upon enzymatic adsorption of TO, DO, MO, and OA respectively.

**Table 4:**
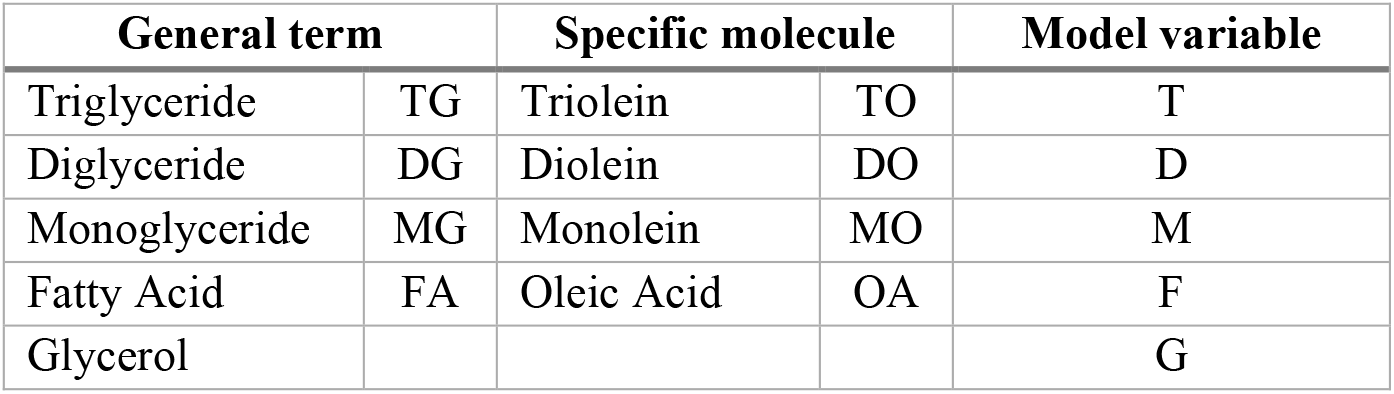
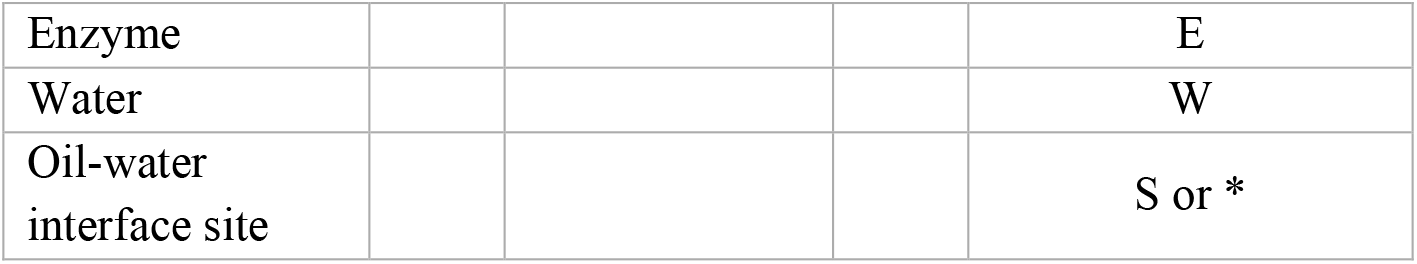
Legend of nomenclature used in the report and in the model for each species

| General term |  | Specific molecule |  | Model variable |
| --- | --- | --- | --- | --- |
| Triglyceride | TG | Triolein | TO | T |
| Diglyceride | DG | Diolein | DO | D |
| Monoglyceride | MG | Monoolein | MO | M |
| Fatty Acid | FA | Oleic Acid | OA | F |
| Glycerol |  |  |  | G |
| Enzyme |  |  |  | E |
| Water |  |  |  | W |
| Oil-water interface site |  |  |  | S or * |

**Figure 3:**
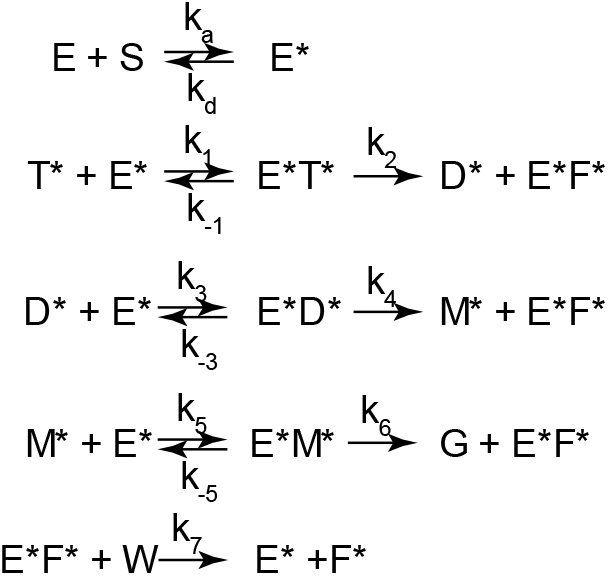
The Ping Pong Bi Bi mechanism of triolein (T*) digestion into diolein (D*), monoolein (M*), glycerol (G), and oleic acid (F*) based on stepwise hydrolysis. S represents the number of sites available at the oil-water interface for enzyme adsorption. The asterisk indicates the molecule is at the oil-water interface. No asterisk indicates the molecule is in solution.

**Figure 4:**
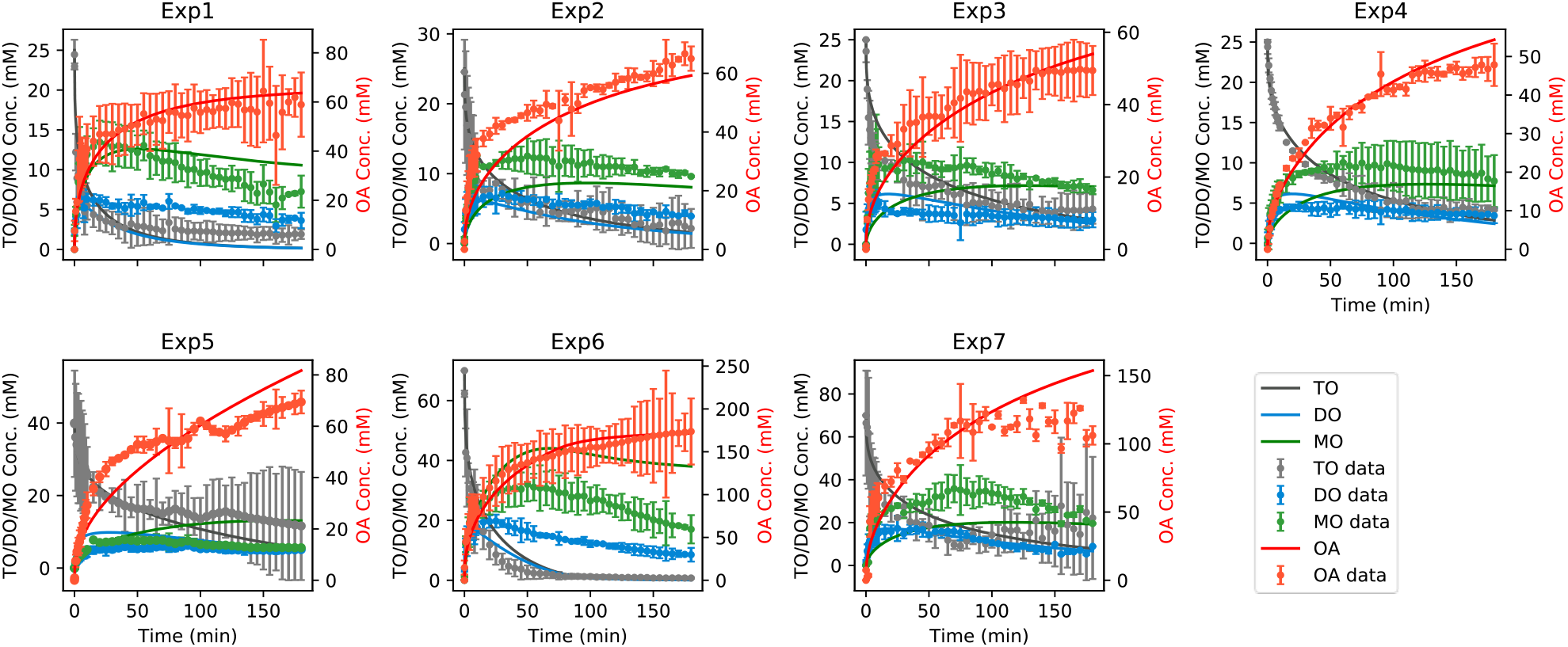
Kinetics of TO digestion, as well as formation and digestion of DO and MO (left axes) and OA formation (right axes), with corresponding lipolysis model fittings. Error bars reflect standard deviation of the mean.

Per the Ping Pong Bi Bi mechanism, the enzyme-fatty acid complex (E*F*) undergoes a further step by reacting with water to produce the fatty acid and “free” (i.e., with an empty active site) adsorbed enzyme (E*) (Paiva et al., 2000). Since DO hydrolysis can result in formation of 2-MO and 1-MO, in the interest of simplicity we did not differentiate between the two isomers and assumed that the interfacial MO follows a similar reaction mechanism to TO and DO.

Enzyme adsorption to the oil-water interface is assumed to follow the Langmuir adsorption model (Hermansyah et al., 2006). Specifically, it is assumed that there is a fixed number of sites (S) at the oil-water interface available for enzyme adsorption per unit surface area. This assumption is particularly important at the reaction limits, such as in the case where the majority of TO and DO have been digested, the interfacial area is small relative to the amount of enzyme available for adsorption and experimental results suggest slow hydrolysis of the remaining TO and DO, in line with the incomplete hydrolysis observed in digestion experiments reported in the literature (Giang et al., 2016; Giang et al., 2015; Li & McClements, 2010). Physiologically, a Langmuir model is also reasonable considering that at such extremities the size of the oil droplets becomes similar to the size of lipase molecules floating in solution. Thus, steric hindrances between the two structures can prevent the concentration dependent adsorption and desorption processes. The forward (adsorption, *r*_*a*_) and reverse (desorption, *r*_*d*_) reaction rates in mol m^-2^ min^-1^ are shown as follows:

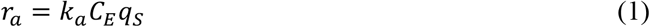

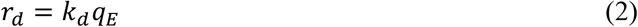

where *k*_*a*_ is the adsorption rate constant in units of m^3^ mol^-1^ min^-1^, *k*_*d*_ is the desorption rate constant in units of min^-1^, *C*_*E*_ is the enzyme concentration in the bulk solution in mol/m^3^, *q*_*E*_ is the concentration of enzyme adsorbed at the oil-water interface in mol/m^2^, and *q*_*S*_ is the enzyme-free number of sites available for adsorption at the oil-water interface in mol/m^2^. The Langmuir model assumes the forward and reverse processes are in constant equilibrium such that *r*_*w*_ = *r*_*d*_. Hence, we define an adsorption coefficient (*K*_*E*_) such that:

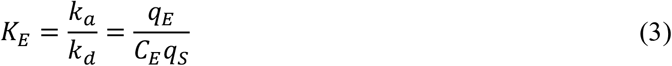

The reaction rates of TO, DO, and MO (*r*_*T*_, *r*_*D*_, and *r*_*M*_) adsorbed at the interface are as follows:

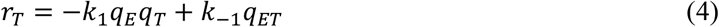

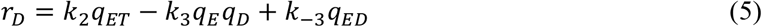

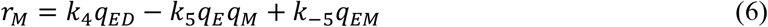

where *q*_*i*_ refers to the interfacial concentration of component *i* in mol/m^2^ oil surface area. The reaction rate of the fatty acid OA (*R*_*F*_ representing an aqueous concentration rate) in solution is as follows:

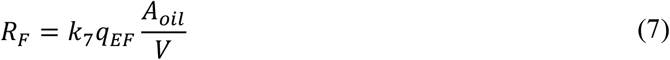

where *V* is the volume of the aqueous phase which is approximately equal to the total solution volume (m^3^), and *A*_*oil*_ is the interfacial area of the oil droplets (m^2^). A pseudo-steady-state assumption (i.e., *dq*_*i*_ /*dt* = 0) for the reaction intermediates, specifically E*T*, E*D*, E*M*, and E*F*, gives the following:

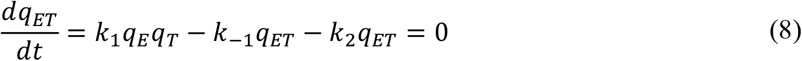

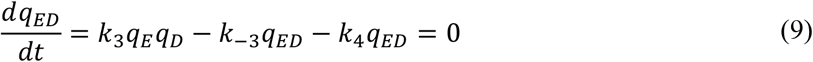

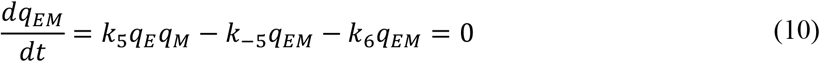

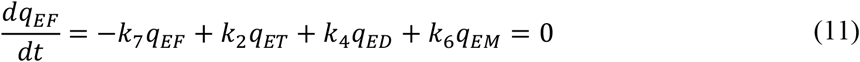

from which the following relationships are obtained:

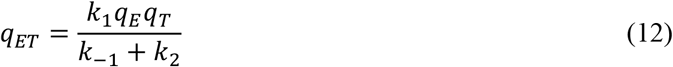

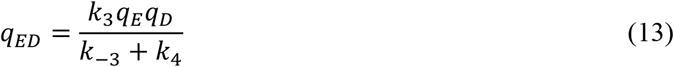

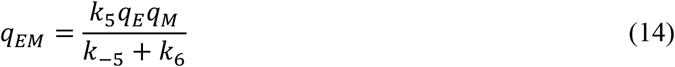

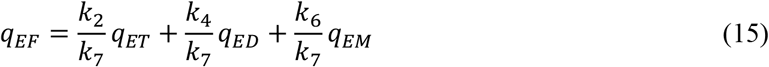

Substituting these relationships into eqns 4 – 7, the following are obtained:

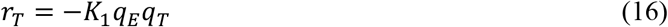

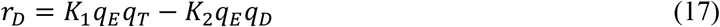

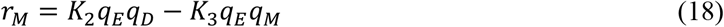

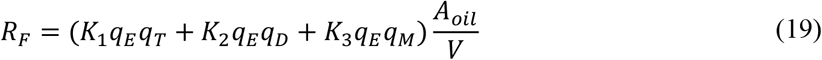

where *K*_1_, *K*_2_, and *K*_3_ are related to the elementary rate constants as shown in Table 5. The rates of lipid digestion can be expressed as change in amount of lipid per unit solution volume due to consumption/generation by hydrolysis as shown in eqns 20 and 21 where *C*_*i*_ refers to the aqueous concentration of component *i* in mol/m^3^:

**Table 5:** Definition of lipolysis coefficients from elementary rate constants

| Coefficients | Relationship to elementary rate constants in Figure 3 |
| --- | --- |
| $K_1$ ($\text{m}^2 \text{mol}^{-1} \text{min}^{-1}$) | $K_1 = \frac{k_1 k_2}{k_{-1} + k_2}$ |
| $K_2$ ($\text{m}^2 \text{mol}^{-1} \text{min}^{-1}$) | $K_2 = \frac{k_3 k_4}{k_{-3} + k_4}$ |
| $K_3 \text{ (m}^2 \text{ mol}^{-1} \text{ min}^{-1}\text{)}$ | $K_3 = \frac{k_5 k_6}{k_{-5} + k_6}$ |
| $K_4 \text{ (m}^2 \text{ mol}^{-1}\text{)}$ | $K_4 = \left(1 + \frac{k_2}{k_7}\right) \frac{k_1}{k_{-1} + k_2}$ |
| $K_5 \text{ (m}^2 \text{ mol}^{-1}\text{)}$ | $K_5 = \left(1 + \frac{k_4}{k_7}\right) \frac{k_3}{k_{-3} + k_4}$ |
| $K_6 \text{ (m}^2 \text{ mol}^{-1}\text{)}$ | $K_6 = \left(1 + \frac{k_6}{k_7}\right) \frac{k_5}{k_{-5} + k_6}$ |
| $K_T \text{ (m)}$ | TO partition coefficient |
| $K_D \text{ (m)}$ | DO partition coefficient |
| $K_M \text{ (m)}$ | MO partition coefficient |
| $K_E \text{ (m}^3 \text{ mol}^{-1}\text{)}$ | $K_E = \frac{k_a}{k_d}$ |
| $q_{E,max} \text{ (mol m}^{-2}\text{)}$ | Maximum interfacial concentration of adsorbed enzyme |

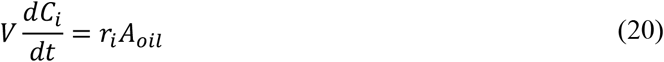

when *i* represents T, D, and M, and

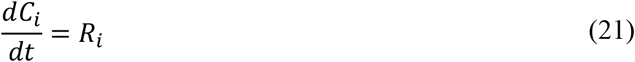

when *i* represents F. Substituting the corresponding reaction rate equations, the overall rates of TO, DO, MO, and OA digestion are obtained as shown:

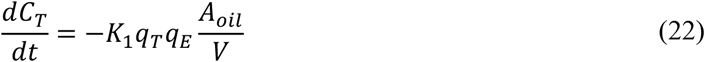

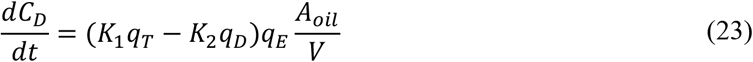

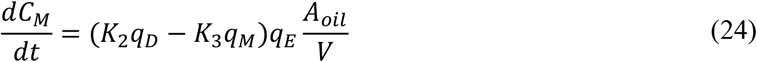

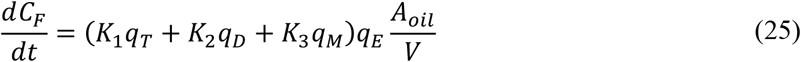

TO and DO partitioning between the interface and the oil core, assumed to be instantaneous as noted above, is reflected in partition coefficients relating corresponding concentrations:

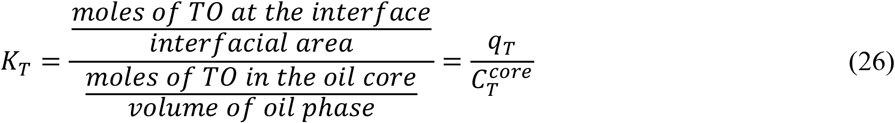

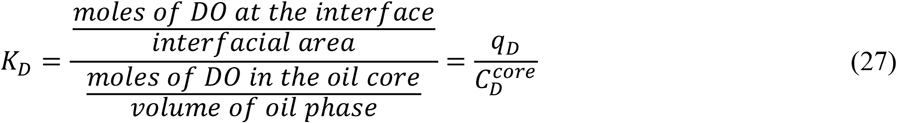

where 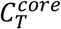 and 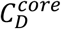 are oil core concentrations of TO and DO, respectively, in units of mol/m^3^ oil volume. Similarly, partitioning of MO and OA between the interface and the aqueous phase is reflected in corresponding partition coefficients (*K*_*M*_ and *K*_*F*_) defined as:

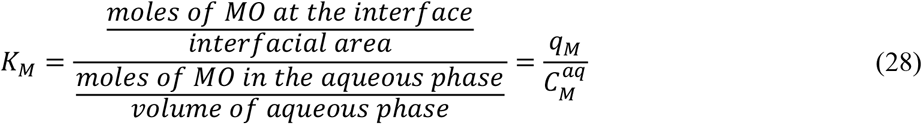

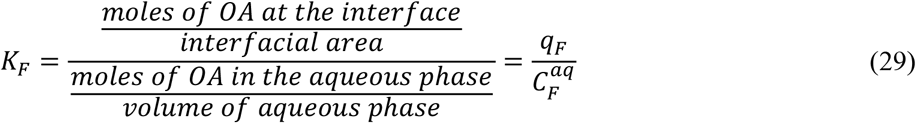

where 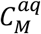 and 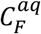 are the concentration of MO and OA in the aqueous phase, respectively, in units of mol/m^3^ solution volume. Using these expressions, interfacial concentrations (*q*_*i*_) can be related to total concentrations (*C*_*i*_) from mass balances as shown:

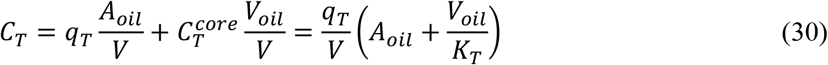

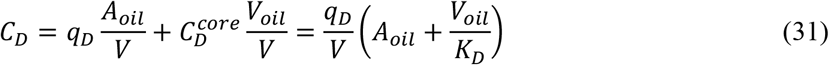

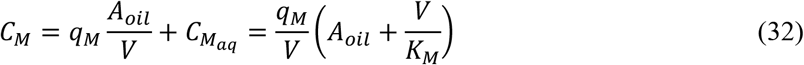

Substituting these relationships between *C*_*i*_ and *q*_*i*_ into eqns 22 – 25, the overall rates of lipid digestion can be expressed as:

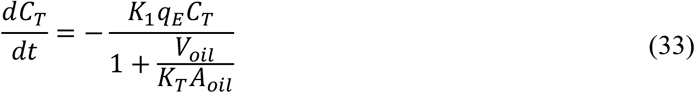

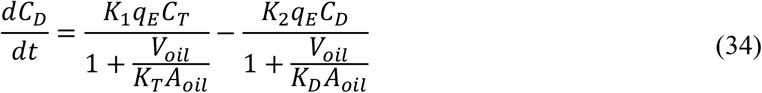

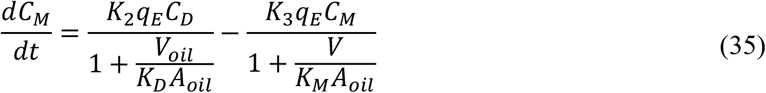

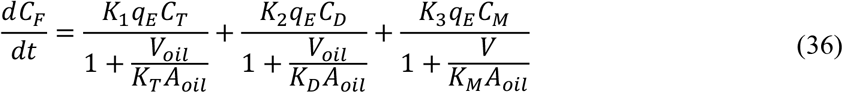

To complete the kinetic model, *q*_*E*_ is related to *C*_*E*_ and to the total enzyme concentration, a known entity, by employing the Langmuir adsorption model and a mass balance on enzyme. First, from a mass balance on the enzyme:

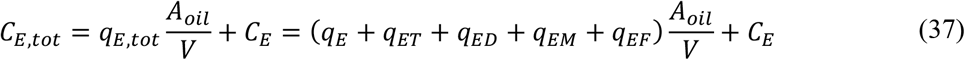

where *C*_*E,tot*_ is the total enzyme concentration. Substituting eqns 12 – 15 into eqn 37, the following is obtained:

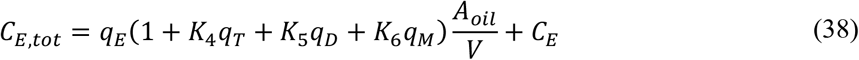

where the coefficients are defined as reported in Table 5. The total number of sites (*q*_*E,max*_) to be considered in the Langmuir adsorption model can be defined as follows:

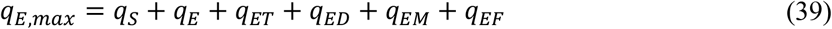

*q*_*S*_ can thus be expressed as follows:

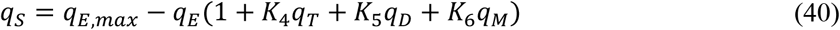

Substituting eqn 40 into eqn 3, the following is obtained:

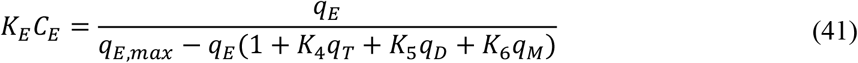

Solving expressions 38 and 41 simultaneously allows determination of *q*_*E*_ as a function of *C*_*E,o*_:

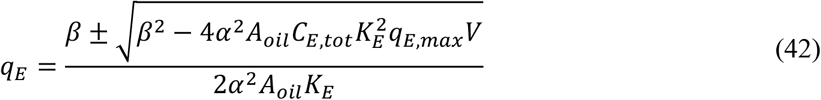

where *α* and *β* are defined as follows:

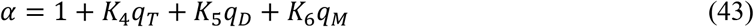

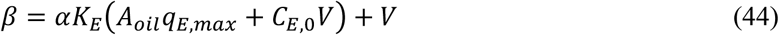

In summary, the lipolysis model has 11 coefficients, specifically *K*_1_, *K*_2_, *K*_3_, *K*_4_, *K*_5_, *K*_6_, *K*_*T*_, *K*_*D*_, *K*_*M*_, *K*_*E*_, and *q*_*E,max*_ (Table 5). It is notable that the fatty acid partition coefficient, *K*_*F*_ (eqn 29), is not needed for the model as the proposed mechanism does not capture interactions between interfacial OA (F*) and enzyme (E*).

A relationship between *A*_*oil*_ and lipid concentrations, specifically *C*_*T*_ and *C*_*D*_, is developed based on the assumption that the number of oil emulsion droplets in solution does not change during lipolysis and is determined prior to the start of digestion based on knowledge of particle size and amount of oil in solution. The equation for quantification of the number of oil droplets in solution (*N*_*oil*_) is as follows:

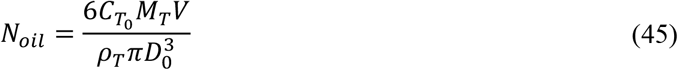

where *D*_*o*_ is the initial average diameter and 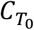 is initial TO concentration. The interfacial area evolution of the oil phase (*A*_*oil*_) was calculated as a function of TO and DO concentrations, as shown in eqn 46.

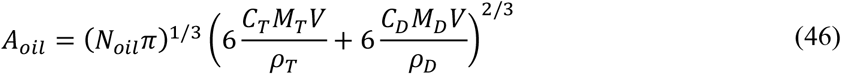

where *C*_*T*_ and *C*_*D*_ represent molar concentrations, *M*_*T*_ and *M*_*D*_ represent molecular weights, and *ρ*_*T*_ and *ρ*_*D*_ represent densities of triolein and diolein, respectively. To fit the kinetic coefficients to the experimental data, a sum of squared normalized differences (SSND) was calculated and minimized using a non-linear regression model. Details of the fitting approach are provided in Supplementary Material.

## 4 Results

### 4.1 Triolein as a representative long chain triglyceride

Triolein was studied as a model long chain triglyceride as its purity and the fact that it is comprised of a single fatty acid chain length enables facile quantification of all products of digestion (i.e., TO, DO, MO, and OA) via HPLC-ELSD and appropriate standards. The same analytical technique is much more challenging with complex natural lipids given the large variety of different types of fatty acids and corresponding monoglycerides, diglycerides, and triglycerides. Triolein digestion kinetics was compared to that of virgin olive oil, linseed oil, palm oil, and coconut oil using a pH-stat autotitrator for quantification of fatty acids produced to assess qualitative similarity and suitability as a model triglyceride. The fatty acid composition and molecular weight of these complex lipids is reported in Table S1 in Supplementary Material. The fatty acid profiles resulting from triglyceride digestion were similar among the long chain triglycerides (Supplementary Figure S1). In contrast, coconut oil, which contains medium chain triglycerides, exhibited faster digestion kinetics (Supplementary Figure S1). These results support suitability of TO as a model long chain triglyceride.

### 4.2 Model fitting to lipolysis training sets and predictions

To enable application of the kinetic model, initial conditions of experiments noted in Table 2 were specified (Table 6). Focusing first on the fed state, the system of equations comprising the proposed model was fit to the corresponding training sets (experiments 1 – 7, Table 6) to obtain the set of kinetic coefficients that resulted in the smallest squared difference between model and experimental results. For fitting, coefficients were allowed to vary within specified ranges set as described in Supplementary Material. Fit kinetic coefficients were then used to compare model predictions to test set experimental lipolysis profiles (experiments 8 – 10, Table 6).

**Table 6:** Initial conditions of lipolysis experiments conducted in this study. Experiments 1 – 10 are conducted in FeSSIF media, whereas experiments 11 – 12 are conducted in FaSSIF media. Calculation of porcine pancreatic lipase (PPL) concentration in mM is described in Supplementary Material.

|  | Exp. no. | Initial TO concentration | Area average diameter (nm) | PPL concentration (mM) | SIF |
| --- | --- | --- | --- | --- | --- |
| **Training Set** | 1 | 25mM | 766 ± 367 | 4.36 x 10<sup>-3</sup> | FeSSIF |
|  | 2 | 25mM | 1567 ± 588 | 4.36 x 10<sup>-3</sup> | FeSSIF |
|  | 3 | 25mM | 2128 ± 1261 | 4.36 x 10<sup>-3</sup> | FeSSIF |
|  | 4 | 25mM | 2049 ± 1294 | 1.45 x 10<sup>-3</sup> | FeSSIF |
|  | 5 | 40mM | 707 ± 265 | 1.45 x 10<sup>-3</sup> | FeSSIF |
|  | 6 | 70mM | 425 ± 113 | 4.36 x 10<sup>-3</sup> | FeSSIF |
|  | 7 | 70mM | 1986 ± 299 | 4.36 x 10<sup>-3</sup> | FeSSIF |
| **Test Set** | 8 | 10mM | 1606 ± 387 | 1.45 x 10<sup>-3</sup> | FeSSIF |
|  | 9 | 20mM | 4076 ± 1328 | 4.36 x 10<sup>-3</sup> | FeSSIF |
|  | 10 | 50mM | 1608 ± 968 | 4.36 x 10<sup>-3</sup> | FeSSIF |
| **Training Set** | 11 | 25mM | 1730 ± 435 | 1.45 x 10<sup>-3</sup> | FaSSIF |
| **Test Set** | 12 | 5mM | 1760 ± 550 | 1.45 x 10<sup>-3</sup> | FaSSIF |

Qualitative examination of experimentally measured lipolysis profiles reveals expected trends (Figure 4) including decreased kinetics with greater emulsion droplet size (experiment 3 vs. 1) and elevated digestion product concentrations with greater starting oil amount (experiment 7 vs. 2), as well as unexpected trends such as lack of difference in digestion kinetics despite a lower enzyme concentration (experiment 4 vs. 3). The convergence results obtained from model SSND calculations using this data indicate that a small number of sets of coefficients can generate similarly good fits (Figure 5, boxed in red and zoomed in on the right panel).

**Figure 5:**
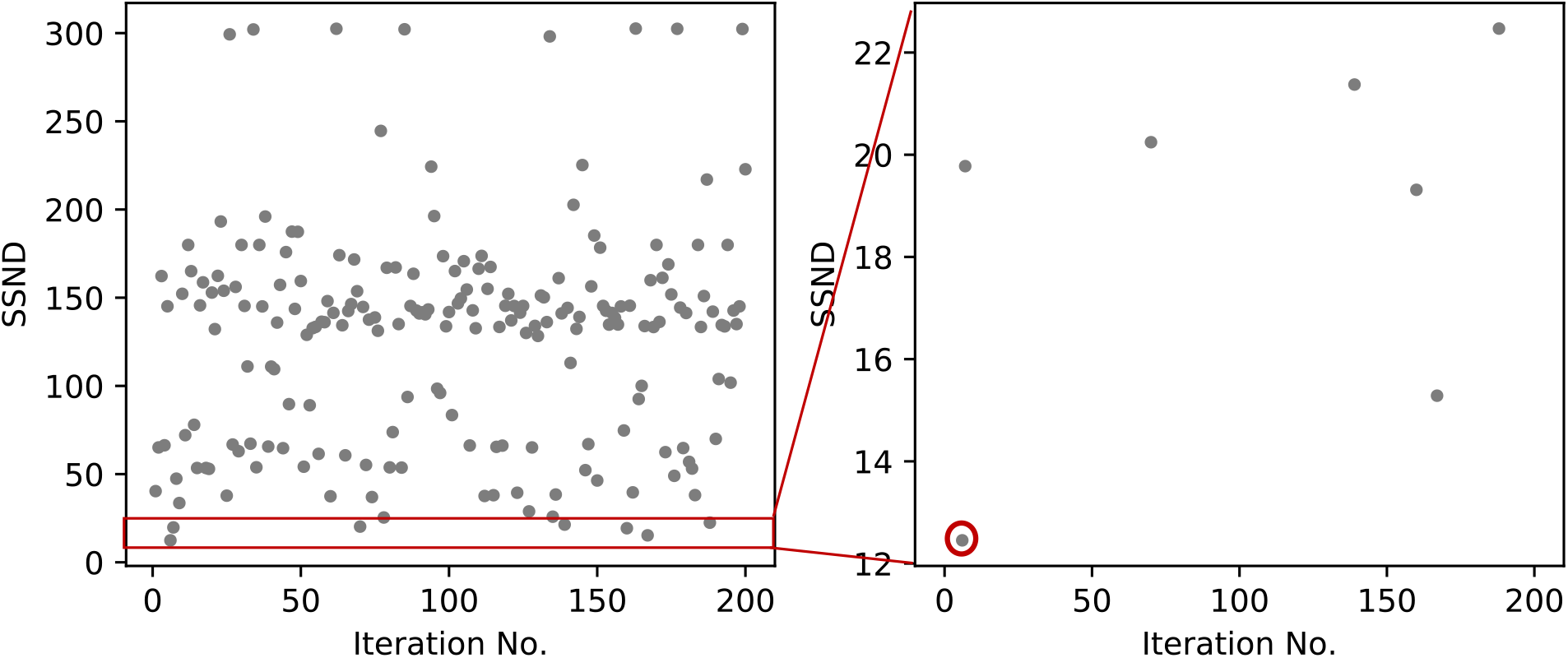
Plot of SSND vs. convergence number of the proposed model in FeSSIF media

Examination of these coefficients (Supplementary Figure S2) reveals that none were pressed against the bounds specified for their values, suggesting that minima could be found in the physiological ranges selected. Additionally, there does not appear to be a correlation among coefficients (e.g., an increase in one causes an increase/decrease in the other), suggesting that the coefficients are linearly independent. The coefficients corresponding to the smallest residual (Table 7) were used to compare model fits to experimental data (Figure 4). The results show reasonable fits overall, with better fits of the TO and OA profiles and less so for profiles of intermediates DO and MO. Effects of increasing droplet size or starting oil amount as well as increased enzyme concentration, as noted above, were reflected in model predictions. The lack of apparent impact of enzyme concentration on lipid profiles in experiment 3 vs. 4 can be explained by examining model predictions of total concentration of enzyme at the interface (*q*_*E,tot*_). Specifically, *q*_*E,tot*_ reaches the Langmuir adsorption model limited maximum interfacial concentration (*q*_*E,max*_) within 30 minutes of commencing lipolysis in experiment 4 and almost immediately in experiment 3 (data not shown), resulting in negligible differences in rates of lipolysis between the two experiments. Sets of coefficients with similar SSND (boxed in Figure 5) produce similarly reasonable predictions of experimental lipolysis profiles (Supplementary Figure S3).

**Table 7:**
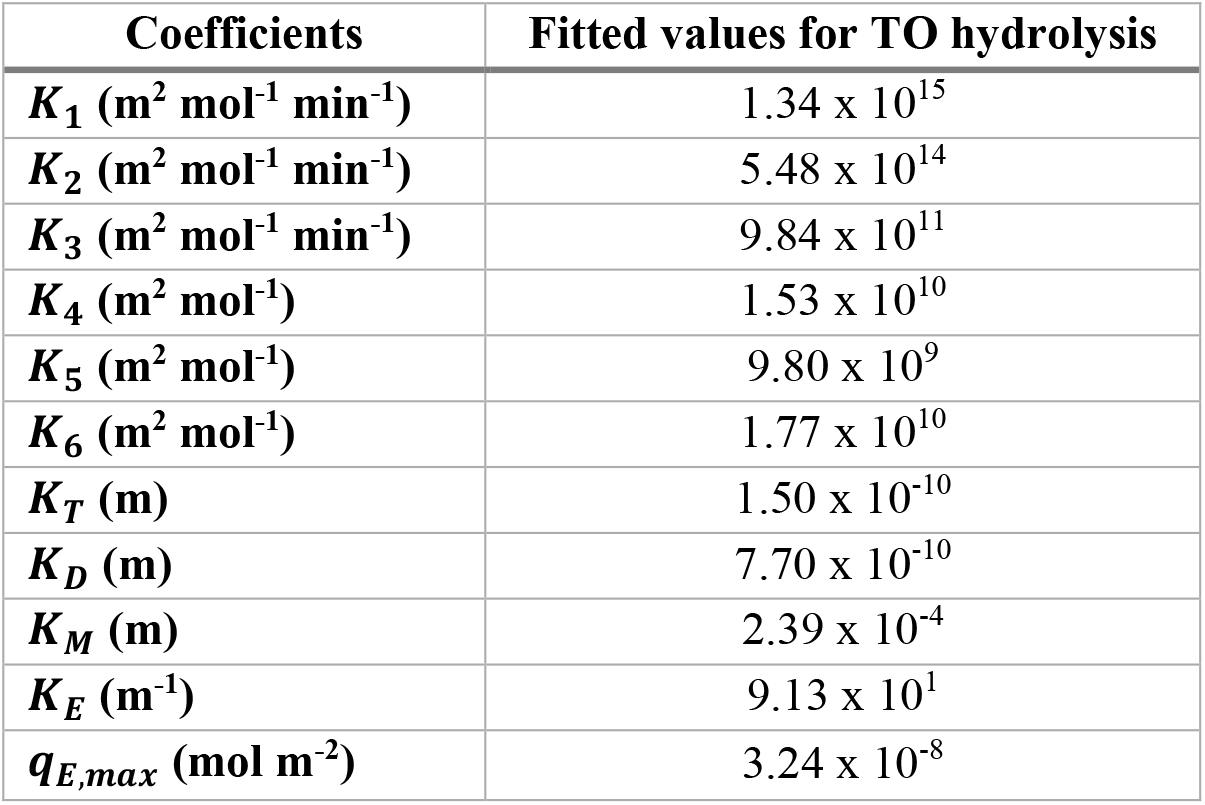
Fitted lipolysis model coefficients for triolein digestion model

| Coefficients | Fitted values for TO hydrolysis |
| --- | --- |
| $K_1$ ($\text{m}^2 \text{mol}^{-1} \text{min}^{-1}$) | $1.34 \times 10^{15}$ |
| $K_2$ ($\text{m}^2 \text{mol}^{-1} \text{min}^{-1}$) | $5.48 \times 10^{14}$ |
| $K_3$ ($\text{m}^2 \text{mol}^{-1} \text{min}^{-1}$) | $9.84 \times 10^{11}$ |
| $K_4$ ($\text{m}^2 \text{mol}^{-1}$) | $1.53 \times 10^{10}$ |
| $K_5$ ($\text{m}^2 \text{mol}^{-1}$) | $9.80 \times 10^9$ |
| $K_6$ ($\text{m}^2 \text{mol}^{-1}$) | $1.77 \times 10^{10}$ |
| $K_T$ (m) | $1.50 \times 10^{-10}$ |
| $K_D$ (m) | $7.70 \times 10^{-10}$ |
| $K_M$ (m) | $2.39 \times 10^{-4}$ |
| $K_E$ ($\text{m}^{-1}$) | $9.13 \times 10^1$ |
| $q_{E,max}$ ($\text{mol m}^{-2}$) | $3.24 \times 10^{-8}$ |

Using the coefficients with lowest SSND (Table 7), the ability of the model to predict experimental fed state lipolysis kinetics from the three test data sets (experiments 8 – 10) was then examined (Figure 6). The results show reasonable comparisons between model predictions and experimental data of the test set.

**Figure 6:**
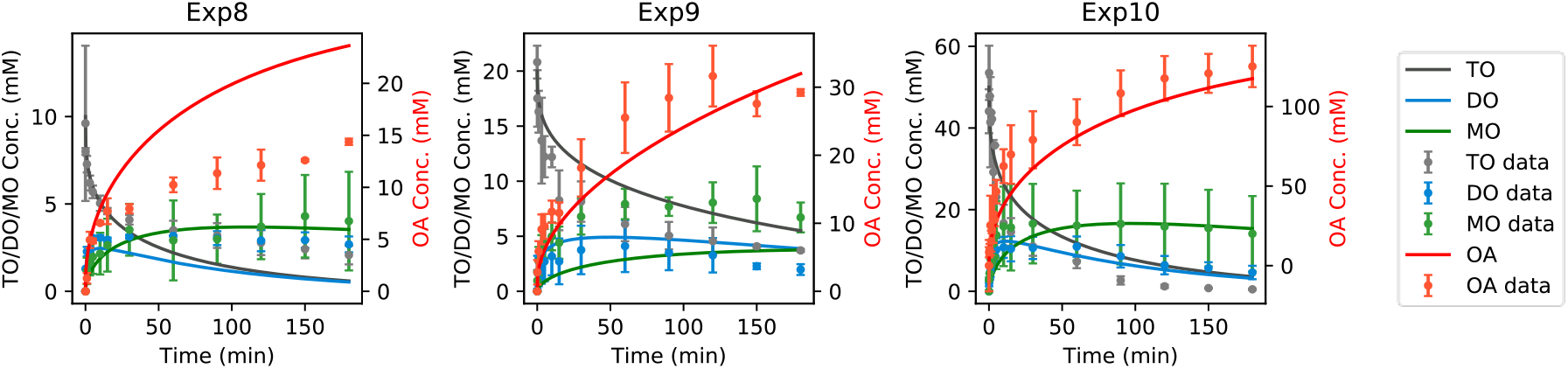
Lipolysis profiles and predictions during digestion of TO from experiments 8 – 10. Error bars reflect standard deviation of the mean (n = 3).

Next, the ability of the model to capture conditions representing the fasted state was examined. While the model does not explicitly take into account concentration of bile components, different model parameters were allowed to vary, reflecting different mechanisms by which bile concentration may impact lipolysis, to fit experimental data obtained using fasted state model bile concentrations (Experiments 11 and 12, Table 6). Allowing *K*_*T*_, *K*_*D*_, and *K*_*M*_ to vary enabled reasonable fits to and predictions of data (Figure S4 and Figure S5 - Supplementary Material), suggesting less interfacial area occupied by bile components in the fasted state may alter distribution of lipid substrates between the emulsion interface and oil core or micelles in solution.

### 4.3 Comparison of model to simpler lipolysis models

While the proposed model is useful in predicting the profiles of all lipid species during digestion in simulated intestinal fluids containing bile micelles, as well as capable of quantifying the distribution of lipid species across phases in the intestinal environment, a major downside is the large number of kinetic coefficients (Table 5) and consequently the large amount of experimental data needed to fit the coefficients and validate the predictive ability of the model. Thus, it was of interest to compare its predictions to those of simpler lipolysis models in the literature proposed by Li & McClements (Li & McClements, 2010) and Buyukozturk et al. (Buyukozturk et al., 2013).

The model by Li & McClements proposed a kinetic expression reflecting the assumption that the rate of FA formation during TG lipolysis is proportional to the interfacial surface area (eqn 47) (Li & McClements, 2010).

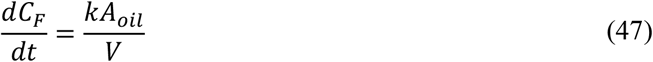

*k* is the kinetic rate constant defined as the number of moles FA produced per unit interfacial area per unit time (mol m^-2^ s^-1^). The authors measured the FA profile via pH-stat titration over 30 min. They observed incomplete digestion in this timeframe and proposed that it was a result of reaction inhibition by the FA produced, which coat the oil-water interface and prevent the enzyme from binding to the oil surface. In the analytical solution of eqn 47, the authors introduced a factor (*φ*_*max*_) representing the extent of digestion to account for the reaction not proceeding to completion. As data reported in our study were obtained over a longer timeframe and show complete depletion of the TG in the 3h of the experiments (experiments 6 and 10) (Figure 4 and Figure 6, respectively), we did not employ the extent factor and used *k* as the only parameter for model fitting. Similarly, to relate *C*_*FA*_ to *A*_*oil*_, the authors assumed that one TG molecule generates two FA molecules. Given that our experimental results show MG depletion during lipolysis, we changed the assumption to indicate that one TG molecule can generate three FA molecules. Following these changes, we fit eqn 47 to the OA profiles from experiment 3 (Figure 7 – dashed lines) and obtained a *k* of 7.50 x 10^−6^ mol m^-2^ min^-1^. This fitted rate constant was then used to predict lipolysis profiles for the other experiments under the same bile and enzyme concentrations (experiments 1, 2, 6, 7, 9, 10) (Figure 7 – dashed lines). Experiment 3 was used for fitting profiles due to relatively low SSND in relation to the other experiments. The profiles of DO and MO are not shown as the model does not capture the kinetics of these components.

**Figure 7:**
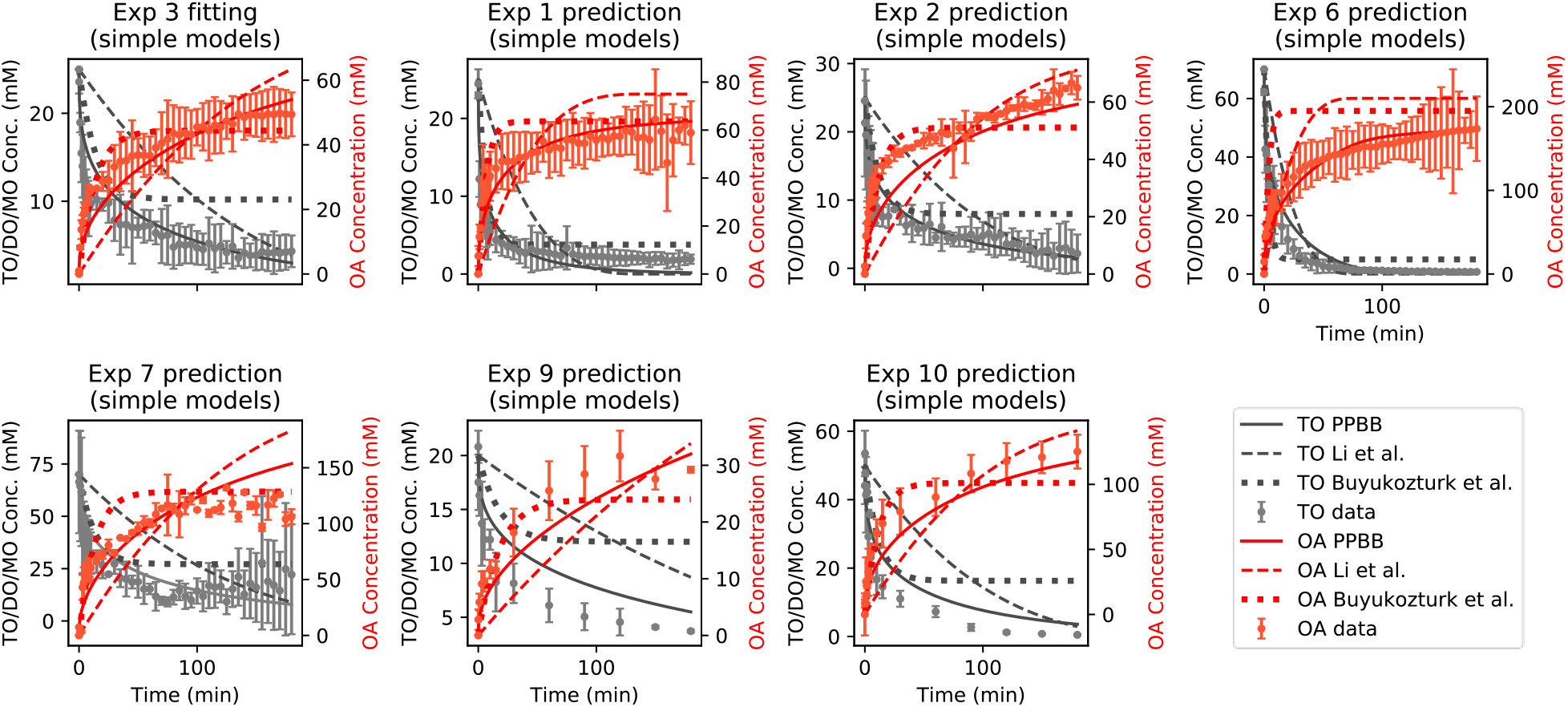
Top left figure - Fitting of lipolysis data from experiment 3 using the Li et al. and Buyukozturk et al. kinetic models. Remaining figures - Comparison of Li et al. and Buyukozturk et al. model predictions to in vitro lipolysis data. All data were obtained in fed state simulated intestinal fluids with fed state enzyme concentration. Data are also compared to the proposed three-step model.

Buyukozturk et al. built a lipolysis model similar to that of Li & McClements where they explicitly introduced an inhibition term proportional to the amount of FA generated over time (eqn 48) (Buyukozturk et al., 2013).

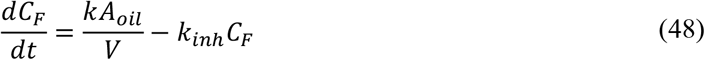

*k*_*inh*_ is the inhibition rate constant due to the FA coating the oil surface (min^-1^). Kinetic rate constants were again determined by fitting eqn 48 to the FA profiles from experiment 3 (Figure 7 – dotted lines). The fitted values for *k* and *k*_*inh*_ were 4.94 x 10^−5^ mol m^2^ min^-1^ and 4.78 x 10^−2^ min^-1^, respectively. Data from experiments 1, 2, 6, 7, 9, and 10 were again compared to predictions of this model (Figure 7 – dotted lines).

The results visibly demonstrate that the simpler mathematical models cannot capture the OA profile during triolein hydrolysis in a simulated intestinal environment as accurately as the proposed model. Specifically, the model by Li and McClements predicts a slow and almost linear depletion of substrate and initial product formation contrary to experimental results. In contrast, the model by Buyukozturk et al. predicts faster hydrolysis at early timepoints and associated fatty acid concentrations which are closer to experimental values, yet it results in an early plateau. The kinetic rate constants of these models were also fit to the other conditions studied experimentally, specifically, digestion in FeSSIF bile micelles + fasted concentrations of pancreatin (Figure S6 – Supplementary Material) and digestion in FaSSIF bile micelles + fasted concentrations of pancreatin (Figure S7 – Supplementary Material). The resulting coefficients are shown in Table S8 – Supplementary Material.

## 5 Discussion

In this study, a modeling framework appropriate for lipid emulsion droplet digestion in the presence of colloidal species, as is carried out routinely in experimental analysis of digestion of lipid-based delivery systems and food lipids, and enabling prediction of reaction intermediates that can markedly impact drug solubility in the intestinal environment, was developed. As the hydrolysis of triglycerides is stepwise and largely dependent on the composition of lipid digestion components at the oil-water interface (Reis et al., 2008), in this study we proposed a three-step kinetic model building upon the Ping Pong Bi Bi mechanism originally developed by Hermansyah et al. (Hermansyah et al., 2006) and adapting the model to account for physiological conditions of the GI tract. Specifically, we accounted for the shrinking of oil emulsion droplets due to triglyceride hydrolysis, partitioning of lipolysis products away from the interface into the oil core or colloidal species in solution, and a modified Langmuir mechanism of enzyme adsorption to the oil-water interface. The kinetic model that was developed generated 11 coefficients which upon fitting to the training set (experiments 1-7), resulted in reasonable comparisons between model predictions and experimental data of the test set (experiments 8-10, Figure 4 and Figure 6). Discrepancies between model predictions and experimental results lie primarily in the fitting of the MO profile which could be due to a variety of factors including coalescence of oil droplets during hydrolysis (Supplementary Material, Figure S8 and Figure S9).

Given some differences between model predictions and experimental data as well as the large number of coefficients, it was of interest to assess the model’s predictive ability compared to simpler models. Predictions of two simpler semi-mechanistic models described by Li & McClements (Li & McClements, 2010) and Buyukozturk et al. (Buyukozturk et al., 2013) were compared to those of the stepwise Ping Pong Bi Bi-based kinetic model proposed in this study (section 3). Specifically, these models result in kinetic expressions requiring fitting of only two kinetic coefficients and consequently require significantly fewer experiments for model fitting. Based on the results shown in Figure 7 and Figures S6 and S7 (Supplementary Material), the simpler models do not accurately predict the experimental profiles relative to the proposed stepwise model. These comparisons indicate that ignoring the stepwise mechanism of FA generation by TG followed by DG followed by MG, is an oversimplification that can dramatically affect the predictive ability of triglyceride lipolysis models. Thus, despite the need for a large number of data points to determine the kinetic coefficients, the ability to capture lipolysis kinetics of all digestion products (i.e., TG, DG, MG, and FA) makes the stepwise Ping Pong Bi Bi model more useful in a variety of applications. In the case of evaluating the utility of lipid-based drug delivery systems in the delivery of lipophilic drug or nutrient compounds, it is valuable to know the composition and disposition of all lipid digestion products in order to understand and quantitatively determine the dynamic effects of triglyceride digestion on overall compound absorption and bioavailability. For example, as both MG and FA partition into micelles, they may both impact the solubilization capacity of the intestinal contents during digestion. As the intestinal solubilization capacity dictates the amount of drug solubilized in the intestinal lumen, it also dictates the driving force for absorption and ultimately bioavailability. Nonetheless, it remains to be seen whether differences in lipolysis kinetic predictions can translate to differences in prediction of absorption and pharmacokinetics of co-dosed oral compounds.

## 6 Conclusions

Lipids are known to impact the delivery of oral compounds (e.g., drugs, nutrients, vitamins), in part by providing an environment into which they can partition, in particular when such compounds have poor aqueous solubility (MacGregor et al., 1997; Persson et al., 2005; Rezhdo et al., 2016; Schawrz et al., 1998). A predictive kinetic model of the fate of lipids in the GI tract upon ingestion capturing the effect of enzyme concentration, the mixed intestinal environment (reflected in emulsion size), and colloidal species (i.e., bile micelles) into which digestion products can partition, could be employed to improve understanding of the impact of lipids on compound absorption. In this study, such a model was developed and applied to experimental data obtained from the hydrolysis of a long chain triglyceride, triolein, by porcine pancreatic lipase in simulated intestinal fluids containing model bile components. The model explicitly considers partitioning of lipid species between the emulsion droplet core, the lipid interface, and mixed bile micelles. It allows determination of quantities of all lipid species present in solution during triglyceride digestion, including monoglyceride and diglyceride. While there is intrinsically a large number of kinetic rate constants due to triglyceride lipolysis being a multi-step kinetic process, there is value in such a model as it can provide guidance on “where” the lipid digestion products are during digestion, i.e., whether they are located at the oil interface, associated with micelles, or inside the oil emulsions. This insight into colloid composition during lipid digestion can offer tremendous value in understanding interactions with compounds co-administered with lipids along the GI tract. For example, lipolysis products that associate with the micelle-rich aqueous phase can increase the solubilization capacity for some co-administered compounds. Understanding and prediction of the quantity of lipolysis products can allow prediction of this solubilization capacity and can thus enable prediction of the impact of dynamic changes in lipid concentration on dissolution kinetics of such compounds. Ultimately, such insight could facilitate rational design of oral delivery vehicles for poorly water-soluble compounds using lipids.

## 7 Data availability

All data is provided in this document. Raw data can be shared upon request.

## 8 Acknowledgements

The authors would like to thank Di Zhu, Christian Gagnon, Samantha Hall, and Tysen DeWaard for collection of some of the data reported in this study. This work was financially supported by the National Institutes of Health (Grant Number R01GM098117) and the National Science Foundation (Grant Number 2015053).

## References

Al-Zuhair, S., Hasan, M., & Ramachandran, K. B. (2003). Kinetics of the enzymatic hydrolysis of palm oil by lipase. Process Biochemistry, 38(8), 1155–1163. 10.1016/S0032-9592(02)00279-0

Al-Zuhair, S., Ramachandran, K. B., & Hasan, M. (2004a). High enzyme concentration model for the kinetics of hydrolysis of oils by lipase. Chemical Engineering Journal, 103(1), 7–11. 10.1016/j.cej.2004.07.001

Al-Zuhair, S., Ramachandran, K. B., & Hasan, M. (2004b). Unsteady-state kinetics of lipolytic hydrolysis of palm oil in a stirred bioreactor. Biochemical Engineering Journal, 19(1), 81–86. 10.1016/j.bej.2003.12.001

Aloulou, A., Puccinelli, D., Sarles, J., Laugier, R., Leblond, Y., & Carriere, F. (2008). In vitro comparative study of three pancreatic enzyme preparations: dissolution profiles, active enzyme release and acid stability. Aliment Pharmacol Ther, 27(3), 283–292. 10.1111/j.1365-2036.2007.03563.x

Amara, S., Bourlieu, C., Humbert, L., Rainteau, D., & Carrière, F. (2019). Variations in gastrointestinal lipases, pH and bile acid levels with food intake, age and diseases: Possible impact on oral lipid-based drug delivery systems. Adv Drug Deliv Rev. 10.1016/j.addr.2019.03.005

Armand, M., Borel, P., Dubois, C., Senft, M., Peyrot, J., Salducci, J., … Lairon, D. (1994). Characterization of emulsions and lipolysis of dietary lipids in the human stomach. Am J Physiol, 266(3 Pt 1), G372–381.

Bakala N’Goma, J. C., Amara, S., Dridi, K., Jannin, V., & Carriere, F. (2012). Understanding the lipid-digestion processes in the GI tract before designing lipid-based drug-delivery systems. Ther Deliv, 3(1), 105–124.

Buyukozturk, F., Di Maio, S., Budil, D. E., & Carrier, R. L. (2013). Effect of ingested lipids on drug dissolution and release with concurrent digestion: a modeling approach. Pharm Res, 30(12), 3131–3144. 10.1007/s11095-013-1238-6

Carriere, F., Barrowman, J. A., Verger, R., & Laugier, R. (1993). Secretion and contribution to lipolysis of gastric and pancreatic lipases during a test meal in humans. Gastroenterology, 105(3), 876–888.

Carrière, F., Renou, C., Lopez, V., de Caro, J., Ferrato, F., Lengsfeld, H., … Verger, R. (2000). The specific activities of human digestive lipases measured from the in vivo and in vitro lipolysis of test meals. Gastroenterology, 119(4), 949–960. 10.1053/gast.2000.18140

Couallier, E., Riaublanc, A., David Briand, E., & Rousseau, B. (2018). Molecular simulation of the water-triolein-oleic acid mixture: Local structure and thermodynamic properties. J Chem Phys, 148(18), 184702. 10.1063/1.5021753

Dahan, A., & Hoffman, A. (2007). The effect of different lipid based formulations on the oral absorption of lipophilic drugs: the ability of in vitro lipolysis and consecutive ex vivo intestinal permeability data to predict in vivo bioavailability in rats. Eur J Pharm Biopharm, 67. 10.1016/j.ejpb.2007.01.017

Di Maio, S., & Carrier, R. L. (2011). Gastrointestinal contents in fasted state and post-lipid ingestion: in vivo measurements and in vitro models for studying oral drug delivery [Comparative Study Review]. J Control Release, 151(2), 110–122.

Fadiloğlu, S., & Söylemez, Z. (1997). Kinetics of lipase-catalyzed hydrolysis of olive oil. Food Research International, 30(3), 171–175. 10.1016/S0963-9969(97)00022-7

Garcia, H. S., Malcata, F. X., Hill, C. G., & Amundson, C. H. (1992). Use of Candida rugosa lipase immobilized in a spiral wound membrane reactor for the hydrolysis of milkfat. Enzyme and Microbial Technology, 14(7), 535–545. 10.1016/0141-0229(92)90124-7

Gargouri, Y., Moreau, H., & Verger, R. (1989). Gastric lipases: biochemical and physiological studies. Biochim Biophys Acta, 1006(3), 255–271.

Gargouri, Y., Pieroni, G., Rivière, C., Lowe, P. A., Saunière, J.-F., Sarda, L., & Verger, R. (1986). Importance of human gastric lipase for intestinal lipolysis: an in vitro study. Biochimica et Biophysica Acta (BBA) - Lipids and Lipid Metabolism, 879(3), 419–423. 10.1016/0005-2760(86)90234-1

Giang, T. M., Gaucel, S., Brestaz, P., Anton, M., Meynier, A., Trelea, I. C., & Le Feunteun, S. (2016). Dynamic modeling of in vitro lipid digestion: Individual fatty acid release and bioaccessibility kinetics. Food Chemistry, 194, 1180–1188. 10.1016/j.foodchem.2015.08.125

Giang, T. M., Le Feunteun, S., Gaucel, S., Brestaz, P., Anton, M., Meynier, A., & Trelea, I. C. (2015). Dynamic modeling highlights the major impact of droplet coalescence on the in vitro digestion kinetics of a whey protein stabilized submicron emulsion. Food Hydrocolloids, 43, 66–72. 10.1016/j.foodhyd.2014.04.037

Goto, M., Goto, M., Nakashio, F., Yoshizuka, K., & Inoue, K. (1992). Hydrolysis of triolein by lipase in a hollow fiber reactor. Journal of Membrane Science, 74(3), 207–214. 10.1016/0376-7388(92)80061-N

Grove, M., Müllertz, A., Nielsen, J. L., & Pedersen, G. P. (2006). Bioavailability of seocalcitol: II: Development and characterisation of self-microemulsifying drug delivery systems (SMEDDS) for oral administration containing medium and long chain triglycerides. European Journal of Pharmaceutical Sciences, 28(3), 233–242. 10.1016/j.ejps.2006.02.005

Hermansyah, H., Kubo, M., Shibasaki-Kitakawa, N., & Yonemoto, T. (2006). Mathematical model for stepwise hydrolysis of triolein using Candida rugosa lipase in biphasic oil–water system. Biochemical Engineering Journal, 31(2), 125–132. 10.1016/j.bej.2006.06.003

Hermansyah, H., Wijanarko, A., Kubo, M., Shibasaki-Kitakawa, N., & Yonemoto, T. (2010). Rigorous kinetic model considering positional specificity of lipase for enzymatic stepwise hydrolysis of triolein in biphasic oil-water system. Bioprocess Biosyst Eng, 33(7), 787–796. 10.1007/s00449-009-0400-3

Ivanova, M., Verger, R., & Panaiotov, I. (1997). Mechanisms underlying the desorption of long-chain lipolytic products by cyclodextrins: application to lipase kinetics in monolayer. Colloids and Surfaces B: Biointerfaces, 10(1), 1–12. 10.1016/S0927-7765(97)00039-8

Kawano, Y., Kawasaki, M., Shiomori, K., Baba, Y., & Hano, T. (1994). Hydrolysis kinetics of olive oil with lipase in a transfer cell. Journal of Fermentation and Bioengineering, 77(3), 283–287. 10.1016/0922-338X(94)90235-6

Kawano, Y., Kiyoyama, S., Shiomori, K., Baba, Y., & Hano, T. (1994). Hydrolysis of olive oil with lipase in a “VibroMixer”. Journal of Fermentation and Bioengineering, 78(4), 293–297. 10.1016/0922-338X(94)90360-3

Labourdenne, S., Ivanova, M. G., Brass, O., Cagna, A., & Verger, R. (1996). Surface behaviour of long-chain lipolytic products (a 1-to-1 mixture of oleic acid and diolein) spread as monomolecular films in the presence of long-chain triglycerides. Colloids and Surfaces B: Biointerfaces, 6(3), 173–180. 10.1016/0927-7765(95)01251-6

Layer, P., & Gröger, G. (1993). Fate of pancreatic enzymes in the human intestinal lumen in health and pancreatic insufficiency. Digestion, 54 Suppl 2, 10–14. 10.1159/000201097

Li, Y., Hu, M., & McClements, D. J. (2011). Factors affecting lipase digestibility of emulsified lipids using an in vitro digestion model: Proposal for a standardised pH-stat method. Food Chemistry, 126(2), 498–505. 10.1016/j.foodchem.2010.11.027

Li, Y., & McClements, D. J. (2010). New Mathematical Model for Interpreting pH-Stat Digestion Profiles: Impact of Lipid Droplet Characteristics on in Vitro Digestibility. Journal of Agricultural and Food Chemistry, 58(13), 8085–8092. 10.1021/jf101325m

MacGregor, K. J., Embleton, J. K., Lacy, J. E., Perry, E. A., Solomon, L. J., Seager, H., & Pouton, C. W. (1997). Influence of lipolysis on drug absorption from the gastro-intestinal tract. Adv Drug Deliv Rev, 25(1), 33–46. 10.1016/S0169-409X(96)00489-9

Maldonado-Valderrama, J., Wilde, P., Macierzanka, A., & Mackie, A. (2011). The role of bile salts in digestion. Adv Colloid Interface Sci, 165(1), 36–46. 10.1016/j.cis.2010.12.002

Mallory, A., Savage, D., Kern, F., Jr., & Smith, J. G. (1973). Patterns of bile acids and microflora in the human small intestine. II. Microflora. Gastroenterology, 64(1), 34–42.

Martinez, O., Wilhelm, A.-M., & Riba, J.-P. (1992). Kinetic study of an enzymatic liquid—liquid reaction: The hydrolysis of tributyrin by Candida cylindracea lipase. Journal of Chemical Technology & Biotechnology, 53(4), 373–378. 10.1002/jctb.280530409

Minekus, M., Alminger, M., Alvito, P., Ballance, S., Bohn, T., Bourlieu, C., … Brodkorb, A. (2014). A standardised static in vitro digestion method suitable for food - an international consensus. Food Funct, 5(6), 1113–1124. 10.1039/c3fo60702j

Mitchell, D. A., Rodriguez, J. A., Carriere, F., Baratti, J., & Krieger, N. (2008). An analytical method for determining relative specificities for sequential reactions catalyzed by the same enzyme: application to the hydrolysis of triacylglycerols by lipases. J Biotechnol, 133(3), 343–350. 10.1016/j.jbiotec.2007.10.012

Mohapatra, S. C., & Hsu, J. T. (1997). Lipase kinetics in organic-water solvent with amphipathic substrate for chiral reaction. Biotechnol Bioeng, 55(2), 399–407. 10.1002/(sici)1097-0290(19970720)55:2<399::Aid-bit17>3.0.Co;2-c

Moreau, H., Gargouri, Y., Lecat, D., Junien, J. L., & Verger, R. (1988). Screening of preduodenal lipases in several mammals. Biochim Biophys Acta, 959(3), 247–252.

Myers, R. A., & Stella, V. J. (1992). Systemic bioavailability of penclomedine (NSC-338720) from oil-in-water emulsions administered intraduodenally to rats. International Journal of Pharmaceutics, 78(1), 217–226. 10.1016/0378-5173(92)90374-B

Olsson, P., Holmback, J., & Herslof, B. (2014). A single step reversed-phase high performance liquid chromatography separation of polar and non-polar lipids. J Chromatogr A, 1369, 105–115. 10.1016/j.chroma.2014.10.010

Paiva, A. L., Balcão, V. M., & Malcata, F. X. (2000). Kinetics and mechanisms of reactions catalyzed by immobilized lipases☆. Enzyme and Microbial Technology, 27(3), 187–204. 10.1016/S0141-0229(00)00206-4

Persson, E. M., Gustafsson, A.-S., Carlsson, A. S., Nilsson, R. G., Knutson, L., Forsell, P., … Abrahamsson, B. (2005). The Effects of Food on the Dissolution of Poorly Soluble Drugs in Human and in Model Small Intestinal Fluids [journal article]. Pharm Res, 22(12), 2141–2151. 10.1007/s11095-005-8192-x

Pilarek, M., & Szewczyk, K. W. (2007). Kinetic model of 1,3-specific triacylglycerols alcoholysis catalyzed by lipases. Journal of Biotechnology, 127(4), 736–744. 10.1016/j.jbiotec.2006.08.012

Plou, F. J., Barandiarán, M., Calvo, M. V., Ballesteros, A., & Pastor, E. (1996). High-yield production of mono- and di-oleylglycerol by lipase-catalyzed hydrolysis of triolein. Enzyme and Microbial Technology, 18(1), 66–71. 10.1016/0141-0229(96)00054-3

Porter, C. J., Kaukonen, A. M., Boyd, B. J., Edwards, G. A., & Charman, W. N. (2004). Susceptibility to lipase-mediated digestion reduces the oral bioavailability of danazol after administration as a medium-chain lipid-based microemulsion formulation. Pharm Res, 21(8), 1405–1412.

Porter, C. J., Kaukonen, A. M., Taillardat-Bertschinger, A., Boyd, B. J., O’Connor, J. M., Edwards, G. A., & Charman, W. N. (2004). Use of in vitro lipid digestion data to explain the in vivo performance of triglyceride-based oral lipid formulations of poorly water-soluble drugs: studies with halofantrine. J Pharm Sci, 93(5), 1110–1121. 10.1002/jps.20039

Reis, P., Holmberg, K., Watzke, H., Leser, M. E., & Miller, R. (2009). Lipases at interfaces: a review. Adv Colloid Interface Sci, 147-148, 237–250. 10.1016/j.cis.2008.06.001

Reis, P., Miller, R., Leser, M., Watzke, H., Fainerman, V. B., & Holmberg, K. (2008). Adsorption of polar lipids at the water-oil interface. Langmuir, 24(11), 5781–5786. 10.1021/la704043g

Rezhdo, O., Di Maio, S., Le, P., Littrell, K. C., Carrier, R. L., & Chen, S.-H. (2017). Characterization of colloidal structures during intestinal lipolysis using small-angle neutron scattering. Journal of Colloid and Interface Science, 499, 189–201. 10.1016/j.jcis.2017.03.109

Rezhdo, O., Speciner, L., & Carrier, R. (2016). Lipid-associated oral delivery: Mechanisms and analysis of oral absorption enhancement. Journal of Controlled Release, 240, 544–560. 10.1016/j.jconrel.2016.07.050

Rice, K. E., Watkins, J., & Hill, C. G., Jr. (1999). Hydrolysis of menhaden oil by a Candida cylindracea lipase immobilized in a hollow-fiber reactor. Biotechnol Bioeng, 63(1), 33–45. 10.1002/(sici)1097-0290(19990405)63:1<33::aid-bit4>3.0.co;2-w

Schawrz, M. A., Raith, K., Dongowski, G., & Neubert, R. H. (1998). Effect on the partition equilibrium of various drugs by the formation of mixed bile salt/phosphatidylcholine/fatty acid micelles. A characterization by micellar affinity capillary electrophoresis. Part IV. J Chromatogr A, 809(1-2), 219–229. 10.1016/s0021-9673(98)00161-7

Shiomori, K., Hayashi, T., Baba, Y., Kawano, Y., & Hano, T. (1995). Hydrolysis rates of olive oil by lipase in a monodispersed OW emulsion system using membrane emulsification. Journal of Fermentation and Bioengineering, 80(6), 552–558. 10.1016/0922-338X(96)87730-0

Tanigaki, M., Sakata, M., Takaya, H., & Mimura, K. (1995). Hydrolysis of palm stearin oil by a thermostable lipase in a draft tube-type reactor. Journal of Fermentation and Bioengineering, 80(4), 340–345. 10.1016/0922-338X(95)94201-2

Tsai, S. W., Wu, G. H., & Chiang, C. L. (1991). Kinetics of enzymatic hydrolysis of olive oil in biphasic organic-aqueous systems. Biotechnol Bioeng, 38(7), 761–766. 10.1002/bit.260380710

van Tilbeurgh, H., Egloff, M. P., Martinez, C., Rugani, N., Verger, R., & Cambillau, C. (1993). Interfacial activation of the lipase-procolipase complex by mixed micelles revealed by X-ray crystallography. Nature, 362(6423), 814–820. 10.1038/362814a0

Verger, R., Mieras, M. C., & de Haas, G. H. (1973). Action of phospholipase A at interfaces. J Biol Chem, 248(11), 4023–4034.

Wang, C. S., Hartsuck, J. A., & Weiser, D. (1985). Kinetics of acylglycerol hydrolysis by human milk lipoprotein lipase. Biochim Biophys Acta, 837(2), 111–118. 10.1016/0005-2760(85)90233-4

Williams, H. D., Anby, M. U., Sassene, P., Kleberg, K., Bakala-N’Goma, J. C., Calderone, M., … Porter, C. J. (2012). Toward the establishment of standardized in vitro tests for lipid-based formulations. 2. The effect of bile salt concentration and drug loading on the performance of type I, II, IIIA, IIIB, and IV formulations during in vitro digestion. Mol Pharm, 9(11), 3286–3300. 10.1021/mp300331z

Williams, H. D., Sassene, P., Kleberg, K., Bakala-N’Goma, J. C., Calderone, M., Jannin, V., … Pouton, C. W. (2012). Toward the establishment of standardized in vitro tests for lipid-based formulations, part 1: method parameterization and comparison of in vitro digestion profiles across a range of representative formulations. J Pharm Sci, 101(9), 3360–3380. 10.1002/jps.23205

Williams, H. D., Sassene, P., Kleberg, K., Calderone, M., Igonin, A., Jule, E., … Porter, C. J. (2013). Toward the establishment of standardized in vitro tests for lipid-based formulations, part 3: understanding supersaturation versus precipitation potential during the in vitro digestion of type I, II, IIIA, IIIB and IV lipid-based formulations. Pharm Res, 30(12), 3059–3076. 10.1007/s11095-013-1038-z

