## Supplementary Material for "Mathematical model of intestinal lipolysis of a long-chain triglyceride"

### 1. Approach to fitting of training set to determine kinetic coefficients

A Backward Differentiation Formula (BDF) method was applied to solve the system of four differential equations using the solve_ivp package in Python 3.7.6 as a function of experimental conditions (Table 2) and kinetic coefficients (Table 5). All experimental data points used for model fitting (experiments 1 – 7 for fed state fitting, experiment 11 for fasted state fitting) were weighted by the maximum possible concentration of species in solution. Specifically, TO, DO, and MO data were normalized by the initial TO concentration whereas OA was normalized by 3 times the initial TO concentration. The best fit was determined using a non-linear least squares regression model to minimize the sum of squared differences between model predictions and experimental data (eqn S1).

$$SSND = \sum_{j}^{7}\sum_{i}^{46}\left(\frac{C_{T_{e_i}} - C_{T_{p_i}}}{C_{T_{0j}}}\right)^2 + \left(\frac{C_{D_{e_i}} - C_{D_{p_i}}}{C_{T_{0j}}}\right)^2 + \left(\frac{C_{M_{e_i}} - C_{M_{p_i}}}{C_{T_{0j}}}\right)^2 + \left(\frac{C_{F_{e_i}} - C_{F_{p_i}}}{3C_{T_{0j}}}\right)^2 \quad \text{(S1)}$$

SSND represents the sum of squared normalized differences. $C_{T_{e_i}}$, $C_{D_{e_i}}$, $C_{M_{e_i}}$, and $C_{F_{e_i}}$ represent the experimentally determined concentrations of the lipid species for each timepoint $i$. Similarly, $C_{T_{p_i}}$, $C_{D_{p_i}}$, $C_{M_{p_i}}$, and $C_{F_{p_i}}$ represent the model predicted concentrations of the lipid species for each timepoint $i$. $C_{T_{0j}}$ represents the initial lipid concentration for experiment $j$. The 7 training sets (i.e., experiments 1 – 7) and 46 timepoints for each training set were used in the calculation and minimization algorithm for SSND. A routine was developed where 100 random initial guesses for the coefficients were chosen within the range specified in section 4 below and were independently allowed to vary such that SSND converges to a minimum. The SSND value from each convergence was recorded. The best set of coefficients was the one that corresponded to the smallest SSND.

### 2. Calculation of concentration of porcine pancreatic lipase

The initial enzyme concentration in solution was calculated based on measured characteristics of pancreatin. The pancreatin employed in this study (P7545 – Sigma Aldrich, St. Louis, MO, US) has been determined to contain 1.1% w/w porcine pancreatic lipase (PPL) (Capolino et al., 2011). From a bicinchoninic acid assay (BCA assay, data not shown), the total protein concentration in the supernatant of the centrifuged pancreatin extract was found to be 65 mg/mL. PPL concentration in the extract is thus 0.7 mg/mL. Upon dilution with the simulated intestinal fluids to reproduce fed conditions at an activity of 2000 U/mL (Minekus et al., 2014) (6 mL of extract in 20mL total volume), as described above, PPL concentration ($C_{E,tot}$) is thus 0.24 mg/mL, which is in agreement with the PPL concentration (0.25 mg/mL) used in *in vitro* studies as reported in the literature (Capolino et al., 2011). Based on a molecular weight of 55 kDa (Capolino et al., 2011), the molar concentration is $C_{E,tot}$ = 4.36 x $10^{-3}$mM. The PPL concentration in the fasted state, when 2 mL pancreatin extract were used in a 20mL total volume, was reduced accordingly.

### 3. Analysis of lipolysis profiles of complex triglycerides vs. triolein

Four "complex" lipids were used in section 4.1 in the main document, specifically, olive oil, linseed oil, palm oil, and coconut oil. Their composition and corresponding molecular weights are reported in Table S1.

*Table S1: Fatty acid composition of different triglycerides used for comparison of digestion kinetics*

| FA \ TG | Olive Oil | Linseed Oil | Palm Oil | Coconut Oil |
|---|---|---|---|---|
| **C18:1** | 81% | 16% | 40% | 6% |
| **C18:2** | 5% | 17% | 10% | 2% |
| **C18:3** | | 58% | | |
| **C18:0** | 3% | 4% | 5% | 3% |
| **C16:0** | 10% | 5% | 44% | 9% |
| **C14:0** | | | | 18% |
| **C12:0** | | | | 48% |
| **C10:0** | | | | 6% |
| **C8:0** | | | | 8% |
| **Reference** | (ThermoScientific, 2022) | (Lewinska et al., 2015) | (Montoya et al., 2014) | (DebMandal & Mandal, 2011) |
| **Molecular weight*** | 876.1 g/mol | 873.3 g/mol | 848.5 g/mol | 659.4 g/mol |

* triglyceride molecular weight was calculated from an average fatty acid molecular weight by adding the molecular weight of a glycerol molecule and subtracting the molecular weight of three water molecules

Quantification of digestion kinetics was conducted via a pH-stat autotitrator as frequently done in the literature (Christiansen et al., 2011; Feeney et al., 2014; Fernandez et al., 2008; Koehl et al., 2020; Tan et al., 2020; Zhu et al., 2013). Measured digestion kinetics were similar among the long chain triglycerides, while coconut oil, which contains medium chain triglycerides, exhibited faster digestion kinetics (Figure S1, left). It can thus be inferred that TO is a representative long chain triglyceride. Since TO digestion could be quantified by both HPLC and titration, results obtained by each technique were compared (Figure S1, right). It is notable that titration underpredicts the fatty acid concentration in solution, particularly at timepoints beyond 30 min. This result is expected considering that the fatty acid concentration is calculated based on the amount of NaOH added to counteract the pH drop caused by the formation of fatty acids. However, not all fatty acids in solution contribute to the pH, as a fraction of the FA in solution are uncharged/protonated and consequently titration and associated calculation of fatty acid concentration does not capture this fraction. To compensate for this effect, back titration is often used, where the pH is adjusted to 9 at the end of the digestion experiment, thus forcing all fatty acids to be charged/unprotonated. Control experiments are also performed to measure the amount of NaOH needed to reach pH 9 in the absence of FA released during digestion. A calculation is then performed to establish a correction factor (Fernandez et al., 2008). However, inherent in this calculation is the assumption that the fraction of uncharged FA is the same irrespective of the total amount of FA in solution. Our data suggest that the underprediction factor is not the same throughout digestion, with the difference becoming greater toward the end of digestion (Figure S1, right). Thus, determination of a correction factor via back titration was not deemed appropriate in our case.

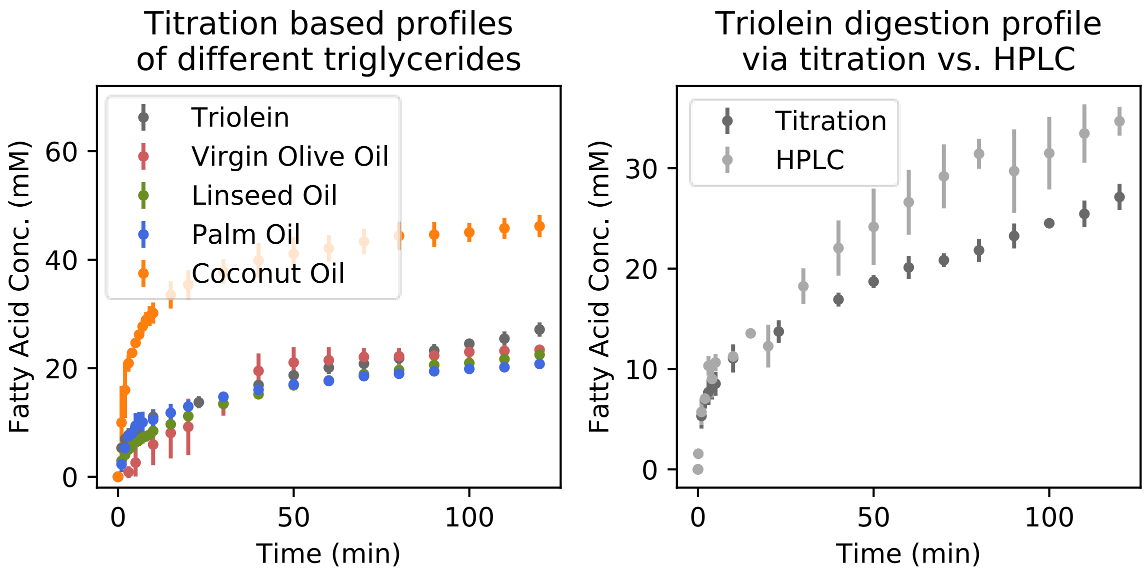


*Figure S1: Left: Fatty acid concentration profiles measured during digestion of different oils (20 mM in FeSSIF media and 6mL pancreatin extract in 20mL total volume). Oils containing mainly long chain triglycerides exhibited similar digestion kinetics, although fatty acid concentrations measured at individual time points were mostly statistically different (statistical analysis not shown due to complexity of the figure). Right: Comparison of fatty acid profiles obtained via titration vs. HPLC analysis following digestion of 20mM triolein in FeSSIF media and 6mL pancreatin extract in 20mL total volume.*

### 4. Determination of physiological ranges of coefficients

To generate physiological values for the coefficients, it was important to define a narrow solution space prior to determining the best fit, i.e., best minimum SSND within that space. Two parameters could be estimated from the literature: $K_T$ and $K_D$. An oil droplet containing pure triglyceride has a fixed ratio of triglyceride at the interface vs. in the oil core assuming there are no amphiphiles coating the interface. The interfacial concentration of triolein is thus dependent on the interfacial area occupied by one TO molecule. A study by Couallier et al. determined the interfacial area of the triolein molecule at a triolein-water interface to be 152.1 Å$^2$ (Couallier et al., 2018). Thus, a surface area of 1 m$^2$ would require 1.092 µmol TO molecules for full coverage (i.e., $q_T$ = 1.092 x 10$^{-6}$ mol/m$^2$). Similarly, the concentration of TO inside an oil core consisting of pure oil can be calculated from the ratio of density (g/mL) and molecular weight (g/mol) of TO. With a density of 0.9078 g/mL and a molecular weight of 885.45 g/mol, the concentration of TO at the oil core ($C_{T_{oil}}$) is 1.025 x 10$^3$ mol/m$^3$. Thus, the maximum possible value for $K_T$ is 1.065 x 10$^{-9}$ m. To set a lower bound, a study by Hamilton et al. was considered, showing that in a triolein droplet stabilized by phosphatidylcholine, 2.8% of the surface contained the pure triglyceride and the rest was covered by the phospholipid (Hamilton & Small, 1981). Thus, we set the lower bound for $K_T$ at 2.8% (i.e., $K_T$ = 2.982 x $10^{-11}$ m). It is likely that in our system surfactants accumulating at the interface can be expelled by the lipase (Reis et al., 2008), thus leaving a fraction larger than 2.8% of the interface available for hydrolysis.

Using the same approach, we could determine the upper and lower bounds for $K_D$. The same study by Couallier et al. determined the interfacial area of one OA molecule to be 36.6 Å$^2$ (Couallier et al., 2018). We estimated the interfacial area of one DO molecule to be equivalent to the difference of the interfacial area of one TO molecule and that of one OA molecule, i.e., 115.5 Å$^2$. Thus, for an oil droplet consisting solely of DO and no amphiphiles at the interface, $q_D$ would be equal to 1/115.5 molecule/Å$^2$ or 1.438 x $10^{-6}$ mol/m$^2$. Similarly, if the oil core consists solely of DO, based on DO density of 0.934 g/mL and molecular weight of 621.11 g/mol, resulting in a DO concentration in the oil core ($C_{D_{oil}}$) of 1.504 x $10^3$ mol/m$^3$. The upper bound for $K_D$ is then determined as the ratio of $q_D$ and $C_{D_{oil}}$ and is equal to 9.561 x $10^{-10}$ m. The lower bound can be estimated at 2.8% of the interfacial area (i.e., $K_D$ = 2.677 x $10^{-11}$ m), although the free fraction of DO at the interface is likely greater than that of TO, given the higher amphiphilicity of DO relative to TO.

For all other coefficients, rough estimations were made. For example, given that $K_1$ dictated the rate of depletion of TO through eqn 33, the slope of the timepoints from the first 3 min of an experiment was used to determine that the order of magnitude (OOM) of $dC_T/dt$ was around $10^0$ mM/min. Assuming $q_T$ to be OOM $10^{-8}$ mol/m$^2$ and given $A_{oil}/V$ OOM of $10^6$ m$^{-1}$, the product $K_1 q_E$ would be about $10^2$. $q_E$ was roughly estimated to be about $10^{-9}$ assuming that all of the 4.36 x $10^{-3}$ mol/m$^3$ of enzyme in solution adheres to the interface and is unbound and available for hydrolysis (i.e., $q_E$ = $C_{E,tot}V/A_{oil}$, where $V/A_{oil}$ = $10^{-6}$ m). Thus, a lower bound for $K_1$ was in the order of $10^{11}$ m$^2$ mol$^{-1}$ min$^{-1}$. The upper bound was initially set to $10^{14}$ m$^2$ mol$^{-1}$ min$^{-1}$ but was gradually increased once convergence results showed the values for $K_1$ constrained against the upper bound. The ranges for $K_2$ and $K_3$ were set to be similar to $K_1$ but broadened slightly depending on the convergence results.

In estimating the upper bound for $q_{E,max}$, we consider that $q_{E,max}$ represents the maximum potential amount of enzyme per surface area that can be adsorbed to the oil-water interface such that hydrolysis kinetics plateaus at that value. As hydrolysis plateaus at the end of digestion (i.e., t = 180 min), we assume at the end of digestion $q_{E,tot}$ is close to $q_{E,max}$. Thus, a limiting case is when we assume that all the enzyme is adsorbed at the oil-water interface such that:

$$q_{E,tot} \approx q_{E,max} \approx \frac{C_{E,tot}V}{A_{oil}|_{180}} \tag{S2}$$

In Table S2 we compute the interfacial area in 20mL solution volume at the end of lipolysis using the following expression:

$$\frac{A_{oil_{180}}}{A_{oil_0}} = \frac{\left(\frac{C_{T_{180}}M_T}{\rho_T} + \frac{C_{D_{180}}M_D}{\rho_D}\right)^{2/3}}{\left(\frac{C_{T_0}M_T}{\rho_T}\right)^{2/3}} \tag{S3}$$

*Table S2: Calculations of interfacial area at start and end of lipolysis (180 min post addition of enzyme in solution) and corresponding $q_{E,max}$ assuming all the enzyme at 180 min is adsorbed at the oil-water interface*

| Exp. no. | Initial TO concentration | Initial interfacial area (m$^2$)* | Final TO/DO concentration | Final interfacial area (m$^2$)** | $q_{E,max}$ (mol/m$^2$)*** |
|---|---|---|---|---|---|
| 1 | 25 mM | 3.664 | 1.38/3.05 mM | 0.991 | 8.79 x 10$^{-8}$ |
| 2 | 25 mM | 1.792 | 1.77/3.30 mM | 0.536 | 1.63 x 10$^{-7}$ |
| 3 | 25 mM | 1.319 | 4.31/3.00 mM | 0.533 | 1.64 x 10$^{-7}$ |
| 4 | 25 mM | 1.370 | 4.18/3.44 mM | 0.563 | 5.16 x 10$^{-8}$ |
| 5 | 40 mM | 2.701 | 11.60/5.09 mM | 1.415 | 2.05 x 10$^{-8}$ |
| 6 | 70 mM | 18.509 | 0.75/8.53 mM | 0.210 | 4.16 x 10$^{-7}$ |
| 7 | 70 mM | 3.958 | 22.16/8.88 mM | 2.169 | 4.02 x 10$^{-8}$ |

*Calculated based on the initial average diameter reported in Table 6

**Calculated from eqn S3

***Calculated from eqn S2

Considering that the values for $q_{E,max}$ represent a maximum, the upper value of $q_{E,max}$ was considered the highest value calculated from experiment 6, i.e., 4 x $10^{-7}$ mol/$m^2$. The minimum value for $q_{E,max}$ was set at 1 x $10^{-12}$ mol/$m^2$.

The range for $K_E$ was estimated assuming that at initial timepoints (t = 0 min) the adsorbed enzyme is unbound to a substrate rather than bound (i.e., $q_E$ is significantly greater than $q_{ET}$, $q_{ED}$, $q_{EM}$, and $q_{EF}$), such that:

$$q_{E,tot}\big|_0 = (q_E + q_{ET} + q_{ED} + q_{EM} + q_{EF})|_0 \approx q_E|_0 \tag{S4}$$

The enzyme mass balance can thus be approximated from eqn 37 as follows:

$$C_{E,tot} \approx q_E \frac{A_{oil}}{V} + C_E \tag{S5}$$

$K_E$ can be determined from the Langmuir model following eqn 3 and a simplification of eqn 39 where the terms representing $q_{ET}$, $q_{ED}$, and $q_{EM}$ are approximated to 0, such that $q_S$ = $q_{E,max}$ - $q_E$. Consequently,

$$K_E = \frac{q_E}{C_E(q_{E,max} - q_E)} \tag{S6}$$

The bounds for $K_E$ are determined assuming that at t = 0, 10% of the enzyme is adsorbed (lower bound) vs. 90% of the enzyme is adsorbed (upper bound). Specifically, assuming 10% is adsorbed to the oil-water interface and 90% of the enzyme is in solution, $C_E$ = 3.92 x $10^{-3}$ mM or 1.31 x $10^{-3}$ mM depending on the targeted enzyme concentration for each experiment. $q_E$ can then be estimated from the initial interfacial area reported in Table S3 and from the mass balance (eqn S5), assuming that $q_E A_{oil}/V = 10\% C_{E,tot}$. Similarly, assuming 10% of the enzyme is in solution and 90% is adsorbed to the oil surface, $C_E$ = 4.36 x $10^{-4}$ mM or 1.45 x $10^{-4}$ mM depending on the target enzyme concentration. Using the upper value for $q_{E,max}$ for each experiment shown in Table S2, corresponding values of $K_E$ can thus be calculated using eqn S6 (Table S3). Based on these values, a range of $10^{-1}$ – $10^4$ $m^3$/mol was estimated for $K_E$.

*Table S3: Estimation of physiological range for $K_E$ using experimental data at the start of digestion*

| Exp. no. | Initial TO concentration | $q_E$ at 10% enzyme adsorbed (mol/m²) | $q_E$ at 90% enzyme adsorbed (mol/m²) | $C_E$ at 10% enzyme adsorbed (mol/m³) | $C_E$ at 90% enzyme adsorbed (mol/m³) | $K_E$ at 10% enzyme adsorbed (m³/mol) | $K_E$ at 90% enzyme adsorbed (m³/mol) |
|---|---|---|---|---|---|---|---|
| **1** | 25 mM | $2.38 \times 10^{-9}$ | $2.14 \times 10^{-8}$ | $3.92 \times 10^{-3}$ | $4.36 \times 10^{-4}$ | 8.3 | 909.5 |
| **2** | 25 mM | $4.87 \times 10^{-9}$ | $2.19 \times 10^{-8}$ | $3.92 \times 10^{-3}$ | $4.36 \times 10^{-4}$ | 9.1 | 1035.6 |
| **3** | 25 mM | $6.61 \times 10^{-9}$ | $4.38 \times 10^{-8}$ | $3.92 \times 10^{-3}$ | $4.36 \times 10^{-4}$ | 11.7 | 1506.8 |
| **4** | 25 mM | $2.12 \times 10^{-9}$ | $5.95 \times 10^{-8}$ | $1.31 \times 10^{-3}$ | $1.45 \times 10^{-4}$ | 36.3 | 4727.2 |
| **5** | 40 mM | $1.07 \times 10^{-9}$ | $9.66 \times 10^{-9}$ | $1.31 \times 10^{-3}$ | $1.45 \times 10^{-4}$ | 45.2 | 6935.0 |
| **6** | 70 mM | $4.71 \times 10^{-10}$ | $4.24 \times 10^{-9}$ | $3.92 \times 10^{-3}$ | $4.36 \times 10^{-4}$ | 0.3 | 23.6 |
| **7** | 70 mM | $2.20 \times 10^{-9}$ | $1.98 \times 10^{-8}$ | $3.92 \times 10^{-3}$ | $4.36 \times 10^{-4}$ | 15.7 | 2519.4 |

$K_M$ was estimated similarly to $K_E$. Specifically, focusing on the end of lipolysis (i.e., at 180 min after the start of digestion), $K_M$ was estimated using eqn 32 assuming anywhere between 10% and 90% of MO is at the oil-water interface (Table S4). Based on these values, a range of $10^{-7}$ – $10^{-3}$ m was proposed for $K_M$.

*Table S4: Estimation of physiological range for $K_M$ using experimental data at 180 min post digestion*

| Exp. no. | Final MO concentration | $C_M^{aq}$ at 10% MO adsorbed (mol/m³) | $C_M^{aq}$ at 90% MO adsorbed (mol/m³) | $q_M$ at 10% MO adsorbed (mol/m²)* | $q_M$ at 90% MO adsorbed (mol/m²)* | $K_M$ at 10% MO adsorbed (m³/mol) | $K_M$ at 90% MO adsorbed (m³/mol) |
|---|---|---|---|---|---|---|---|
| **1** | 6.35 mM | 5.72 | 0.64 | $1.28 \times 10^{-5}$ | $1.15 \times 10^{-4}$ | $2.24 \times 10^{-6}$ | $1.82 \times 10^{-4}$ |
| **2** | 8.11 mM | 7.30 | 0.81 | $3.03 \times 10^{-5}$ | $2.72 \times 10^{-4}$ | $4.15 \times 10^{-6}$ | $3.36 \times 10^{-4}$ |
| **3** | 6.62 mM | 5.96 | 0.66 | $2.48 \times 10^{-5}$ | $2.24 \times 10^{-4}$ | $4.17 \times 10^{-6}$ | $3.38 \times 10^{-4}$ |
| **4** | 7.73 mM | 6.96 | 0.77 | $2.74 \times 10^{-5}$ | $2.47 \times 10^{-4}$ | $3.95 \times 10^{-6}$ | $3.20 \times 10^{-4}$ |
| **5** | 8.19 mM | 7.37 | 0.82 | $4.89 \times 10^{-6}$ | $4.40 \times 10^{-5}$ | $6.63 \times 10^{-7}$ | $5.37 \times 10^{-5}$ |
| **6** | 17.02 mM | 15.32 | 1.70 | $1.62 \times 10^{-4}$ | $1.46 \times 10^{-3}$ | $1.06 \times 10^{-5}$ | $8.58 \times 10^{-4}$ |
| **7** | 19.59 mM | 17.63 | 1.96 | $1.81 \times 10^{-5}$ | $1.96 \times 10^{-4}$ | $1.02 \times 10^{-6}$ | $8.30 \times 10^{-5}$ |

*Calculated using the final interfacial area reported in Table S2

$K_4$, $K_5$, and $K_6$ were approximated based on the first term in the denominator of eqn 41. Specifically, as $q_T$ and $q_D$ are in range of $10^{-8}$ or lower ($q_M$ is assumed to be in a similar range), $K_4$, $K_5$, and $K_6$ must be in the range of $10^{8}$ in order for the product $K_4 q_T$, $K_5 q_D$, and $K_6 q_M$ to be in a similar range as the first term (i.e., 1). Physiologically, each term in the denominator when multiplied by $q_E$ represents the interfacial concentration of substrate-bound enzyme ($q_{ET}$, $q_{ED}$, $q_{EM}$, $q_{EF}$) or substrate unbound enzyme ($q_E$). If one or all of the products $K_4 q_T$, $K_5 q_D$, or $K_6 q_M$ are significantly greater than 1, it indicates that the adsorbed enzyme is highly unlikely to be in its free form (i.e., E*). Instead, it is mainly bound to a substrate (e.g., E*T*, E*D*, or E*M*). Alternatively, if the values of one or all of the products $K_4 q_T$, $K_5 q_D$, or $K_6 q_M$ are significantly smaller than 1, the model favors the adsorbed enzyme to be primarily present in its free from (i.e., E*) rather than bound to a substrate. Thus, to provide this flexibility for the model, $K_4$, $K_5$, and $K_6$ were set to vary between 1 x $10^6$ – 1 x $10^{12}$ $m^2$/mol.

Considering that some of these ranges were determined from rough assumptions, the regression model was allowed to randomly select initial guesses from the parameter range specified above. Further, the bounds included in the least-squares regression model were expanded beyond the estimated parameter range in an effort to scrutinize the likelihood of our assumptions. For example, $K_E$ was determined based on the assumption that between 10% - 90% of the enzyme adsorbs to the oil-water interface. The assumption could also have been between 1% to 99%, which would lead to a 10-fold smaller value for the lower bound and 10-fold larger value for the upper bound for $K_E$. The selected range of parameters is shown in Table S5.

*Table S5: Parameter range used for model fitting*

| Constants | Parameter range for initial guess | Bounds for least squares regression |
|---|---|---|
| $K_1$ ($m^2$ $mol^{-1}$ $min^{-1}$) | 1 x $10^{11}$ – 1 x $10^{16}$ | 1 x $10^8$ – 1 x $10^{17}$ |
| $K_2$ ($m^2$ $mol^{-1}$ $min^{-1}$) | 1 x $10^{11}$ – 1 x $10^{17}$ | 1 x $10^8$ – 1 x $10^{17}$ |
| $K_3$ ($m^2$ $mol^{-1}$ $min^{-1}$) | 1 x $10^{10}$ – 1 x $10^{15}$ | 1 x $10^8$ – 1 x $10^{17}$ |
| $K_4$ ($m^2$ $mol^{-1}$) | 1 x $10^6$ – 1 x $10^{12}$ | 1 x $10^5$ – 1 x $10^{15}$ |
| $K_5$ ($m^2$ $mol^{-1}$) | 1 x $10^6$ – 1 x $10^{12}$ | 1 x $10^5$ – 1 x $10^{15}$ |
| $K_6$ ($m^2$ $mol^{-1}$) | 1 x $10^6$ – 1 x $10^{12}$ | 1 x $10^5$ – 1 x $10^{15}$ |
| $K_T$ (m) | 2.984 x $10^{-11}$ – 1.065 x $10^{-9}$ | 2.984 x $10^{-11}$ – 1.065 x $10^{-9}$ |
| $K_D$ (m) | 2.677 x $10^{-11}$ – 9.561 x $10^{-10}$ | 2.677 x $10^{-11}$ – 9.561 x $10^{-10}$ |
| $K_M$ (m) | 1 x $10^{-10}$ – 1 x $10^1$ | 1 x $10^{-10}$ – 1 x $10^1$ |
| $K_E$ ($m^3$ $mol^{-1}$) | 1 x $10^{-1}$ – 1 x $10^4$ | 1 x $10^{-2}$ – 1 x $10^6$ |
| $q_{E,max}$ (mol $m^{-2}$) | 1 x $10^{-12}$ – 4 x $10^{-7}$ | 1 x $10^{-13}$ – 1 x $10^{-5}$ |

### 5. Analysis of model parameter and predictions resulting from lowest SSND fits

The coefficients from the top 7 fits reported in Figure 4 and their relation to upper and lower bounds set for the solution space are shown in Figure S2.

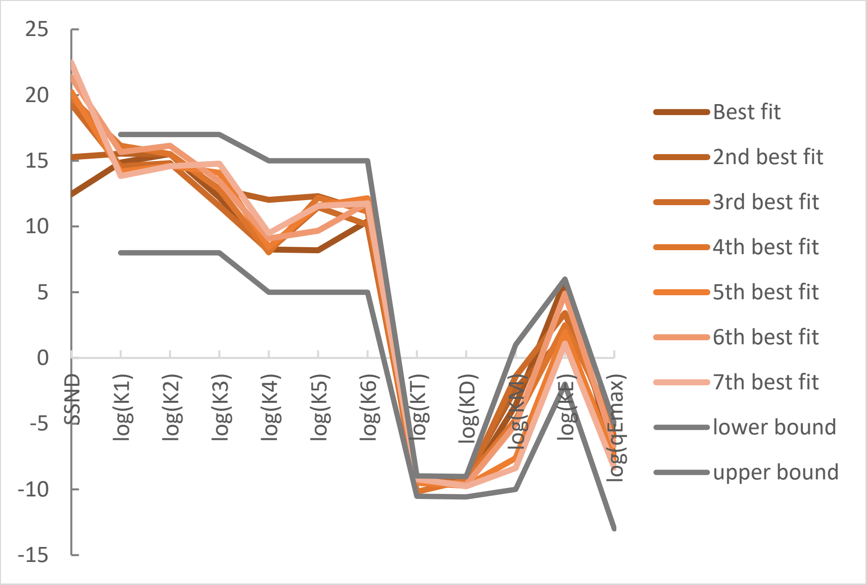


*Figure S2: Coefficient values corresponding to the best fits, i.e., to the lowest SSND values in ascending order based on the value of SSND*

In addition to the results corresponding to the best fit presented in the main document, it was of interest to understand how different a SSND of 12.5 (best fit) is from a SSND of 15 ($2^{nd}$ best fit) and a SSND of 20 ($3^{rd}$ – $7^{th}$ best fits) in terms of ability to capture the profiles of each lipid species. Thus, we used each of the 7 sets of coefficients to predict the lipolysis profile of experiment 4 as the profile with the lowest SSND (Figure S3). The results show that all fits with the exception of the $3^{rd}$ fit are quite similar, suggesting that all 7 sets of fitted coefficients could reasonably capture the rates of hydrolysis of TO and its digestion products under different initial conditions.

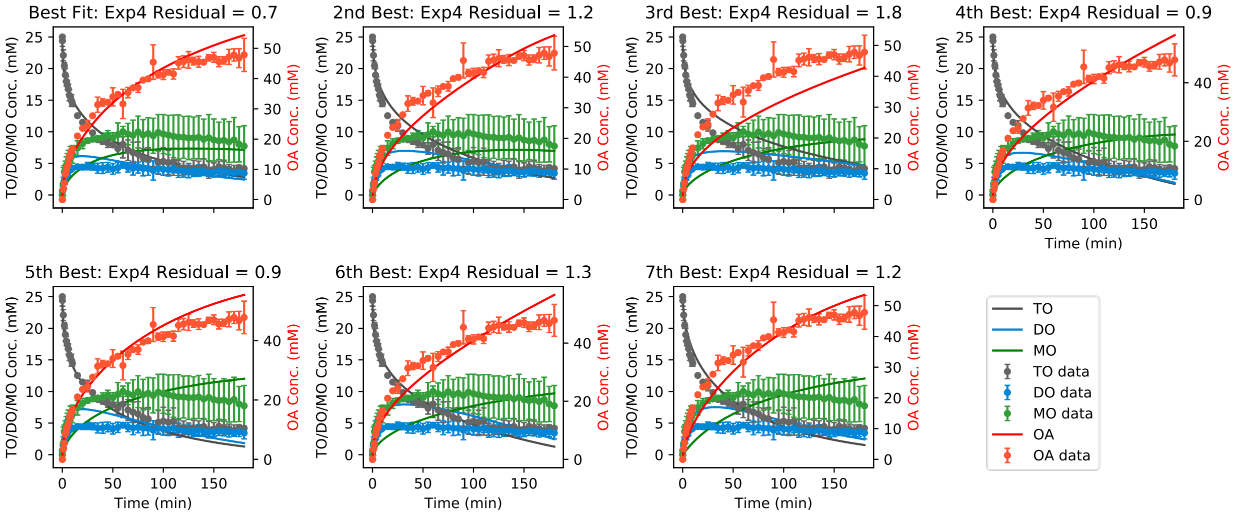


*Figure S3: Comparison of model predictions based on the top seven best fitted coefficient sets to experimental TO lipolysis profiles (experiment 4).*

### 6. Tentative models for reflecting impact of bile concentration on lipolysis

While bile salts are known to mediate lipase binding to the oil-water interface (Brockman, 2000), it is not known whether the enzyme binding efficiency is dependent on the concentration of bile. It is also not known how the number of micelles in the bulk solution impacts the capacity of micelles and aqueous environment to "carry" amphiphilic lipolysis products, such as MG and FA. In the absence of such information, the proposed model does not explicitly incorporate the effect of bile concentration on lipolysis kinetics. However, it can be used as a tool to evaluate these mechanisms. Experiments 4 and 11 have similar initial TO concentration, emulsion droplet size, and enzyme concentration (Table 6). However, they differ in the amount of bile in solution (Table 6). Comparing the lipid profiles, experiment 11 (Figure S4) shows a faster rate of hydrolysis than experiment 4 (Figure 4), suggesting that the higher concentration of bile components has some inhibitory effect on digestion. To predict lipolysis kinetics in FaSSIF media, we allowed specific parameters to vary reflecting potential mechanisms of bile impact, as described in Table S6:

*Table S6: Potential models describing lipolysis in the fasted state*

| Model No. | Proposed mechanism of bile effect on lipolysis | Coefficients to vary |
|---|---|---|
| 1 | Assuming bile concentration has no effect on lipolysis | None |
| 2 | Assuming the ratio of NaTDC at the oil surface vs. in the bulk<br>solution is the same in FaSSIF and FeSSIF for the same amount | $q_{E,max}$ |

| | | |
|---|---|---|
| | of oil and droplet size, in FaSSIF there should be fewer NaTDC molecules per unit interfacial area. Thus, it could be hypothesized that there is a larger number of active sites at the oil-water interface for the enzyme to adsorb to. Consequently, digestion occurs more quickly in the fasted state. $q_{E,max}$ was re-fit. | |
| 3 | In addition to an altered number of active sites per unit interfacial area, the different amount of NaTDC at the interface could affect $K_E$, given that NaTDC is known to mediate colipase adsorption to the oil-water interface. $q_{E,max}$ and $K_E$ were re-fit. | $q_{E,max}$, $K_E$ |
| 4 | As the amount of NaTDC at the oil-water interface is diminished in the fasted state, it is likely that the substrates (i.e., TO, DO, MO) can be found in greater quantities at the interface relative to at the oil core (i.e., TO, DO) or in solution (i.e., MO). More substrate available at the interface results in faster digestion kinetics. $K_T$, $K_D$, and $K_M$ were re-fit. | $K_T$, $K_D$, $K_M$ |
| 5 | A combination of the last three mechanisms | $q_{E,max}$, $K_E$, $K_T$, $K_D$, $K_M$ |

The best fit from each situation described in Table S6 and the associated SSND as an indication of the proposed mechanism ability to capture experimental data is reported in Figure S4. The results indicate that mechanisms 4 and 5 are the most likely as they result in the smallest residual.

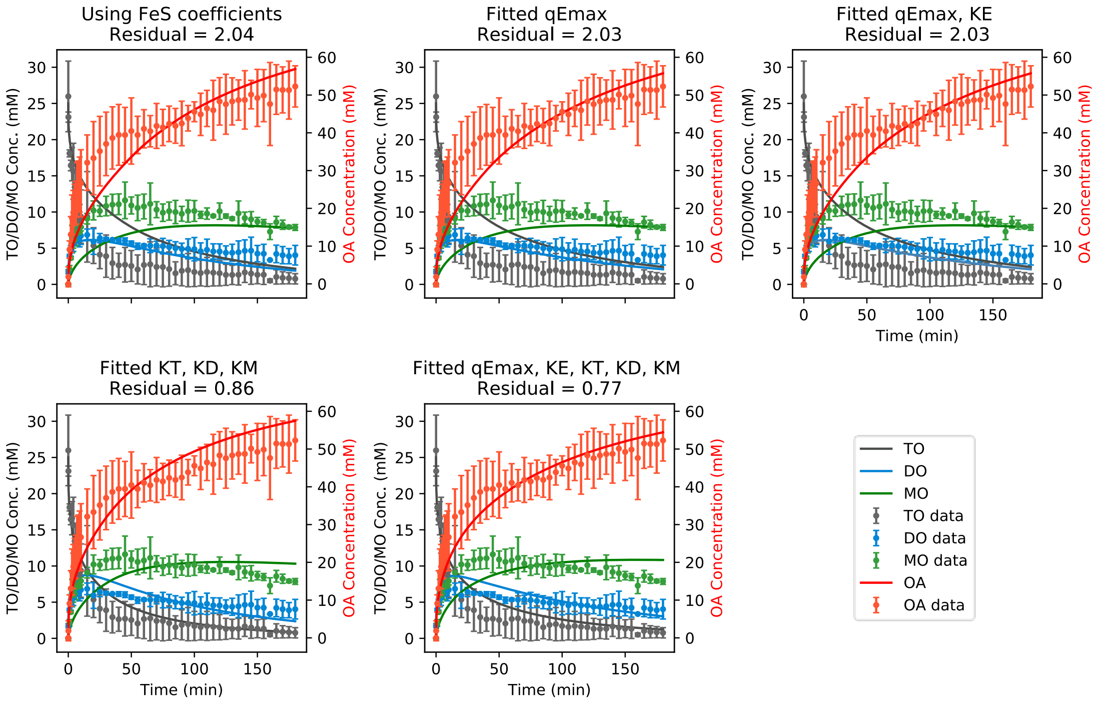


*Figure S4: Fits of experiment 11 using different models of the impact of lower bile component levels in fasted state as reported in Table S6*

The fitted coefficients and goodness of fit for each model are reported in Table S7.

*Table S7: Values of coefficients during digestion in fasted state simulated intestinal fluids*

| Constants | FeSSIF fitted coefficients | Fitted $q_{E,max}$ | Fitted $q_{E,max}$ and $K_E$ | Fitted $K_T$, $K_D$, and $K_M$ | Fitted $q_{E,max}$, $K_E$, $K_T$, $K_D$, $K_M$ |
|---|---|---|---|---|---|
| $K_1$ (m$^2$ mol$^{-1}$ min$^{-1}$) | 1.34 x $10^{15}$ | Same as FeSSIF | Same as FeSSIF | Same as FeSSIF | Same as FeSSIF |
| $K_2$ (m$^2$ mol$^{-1}$ min$^{-1}$) | 5.48 x $10^{14}$ | Same as FeSSIF | Same as FeSSIF | Same as FeSSIF | Same as FeSSIF |
| $K_3$ (m$^2$ mol$^{-1}$ min$^{-1}$) | 9.84 x $10^{11}$ | Same as FeSSIF | Same as FeSSIF | Same as FeSSIF | Same as FeSSIF |
| $K_4$ (m$^2$ mol$^{-1}$) | 1.53 x $10^{10}$ | Same as FeSSIF | Same as FeSSIF | Same as FeSSIF | Same as FeSSIF |
| $K_5$ (m$^2$ mol$^{-1}$) | 9.80 x $10^{9}$ | Same as FeSSIF | Same as FeSSIF | Same as FeSSIF | Same as FeSSIF |
| $K_6$ (m$^2$ mol$^{-1}$) | 1.77 x $10^{10}$ | Same as FeSSIF | Same as FeSSIF | Same as FeSSIF | Same as FeSSIF |
| $K_T$ (m) | 1.50 x $10^{-10}$ | Same as FeSSIF | Same as FeSSIF | 9.42 x $10^{-10}$ | 9.79 x $10^{-10}$ |

| | | | | | |
|---|---|---|---|---|---|
| $K_D$ (m) | 7.70 x $10^{-10}$ | Same as FeSSIF | Same as FeSSIF | 2.32 x $10^{-10}$ | 2.46 x $10^{-10}$ |
| $K_M$ (m) | 2.39 x $10^{-4}$ | Same as FeSSIF | Same as FeSSIF | 8.21 x $10^{-2}$ | 8.27 x $10^{-3}$ |
| $K_E$ ($m^{-1}$) | 9.13 x $10^{1}$ | Same as FeSSIF | 5.96 x $10^{4}$ | Same as FeSSIF | 1.39 x $10^{4}$ |
| $q_{E,max}$ (mol $m^{-2}$) | 3.24 x $10^{-8}$ | 2.75 x $10^{-8}$ | 2.75 x $10^{-8}$ | Same as FeSSIF | 2.29 x $10^{-8}$ |
| **Residual (SSND)** | 2.04 | 2.03 | 2.03 | 0.86 | 0.77 |

We selected model 4 which fit $K_T$, $K_D$, and $K_M$ considering that re-fitting these three coefficients had the largest impact in diminishing the residual while also keeping a low number of coefficients to vary. Using this set of coefficients from Table S7, we compared model predictions in the fasted state to experiment 12 data (Figure S5). Indeed, model 4, considering a potential impact of bile components on partitioning of TO, DO, or MO at the oil interface, offers a reasonable prediction of TO digestion kinetics in the fasted state.

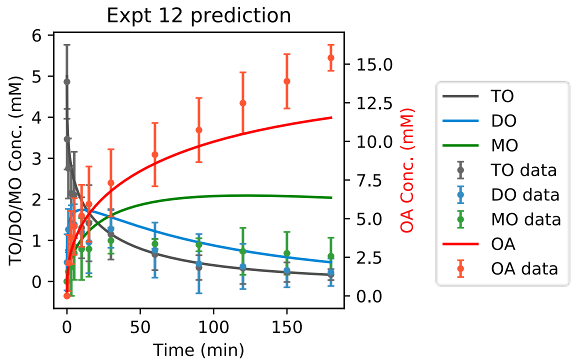


*Figure S5: Lipid profile based on model predictions using coefficients in Table S7 compared to experimental data from experiment 12.*

### 7. Comparisons of proposed model to simpler lipolysis models using data obtained with fasted state enzyme concentration

In addition to model fitting and predictions reported in the main document (Figure 7), the kinetic rate constants of the models by Li et al. and Buyukozturk et al. were obtained also in FeSSIF + fasted concentrations of pancreatin (Figure S6) and FaSSIF + fasted concentrations of pancreatin (Figure S7). The resulting coefficients are summarized in Table S8.

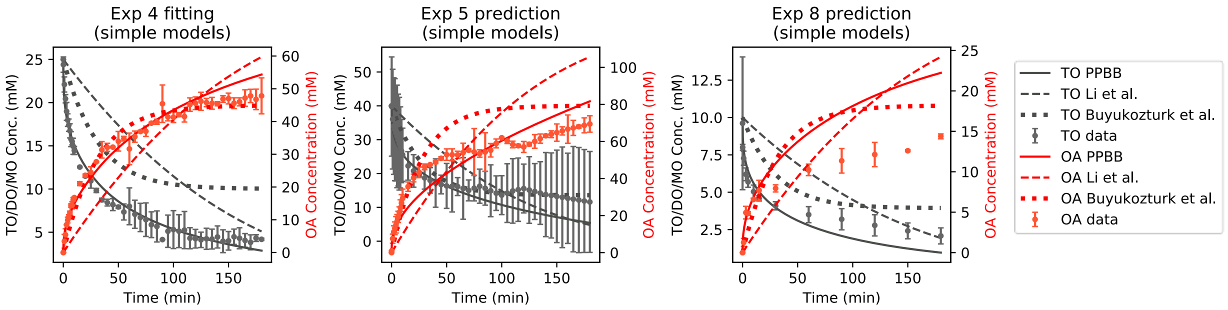


*Figure S6: Left - Fitting of lipolysis data from experiment 4 to models by Li et al. and Buyukozturk et al.; Right – Comparison of model predictions among the three models.*

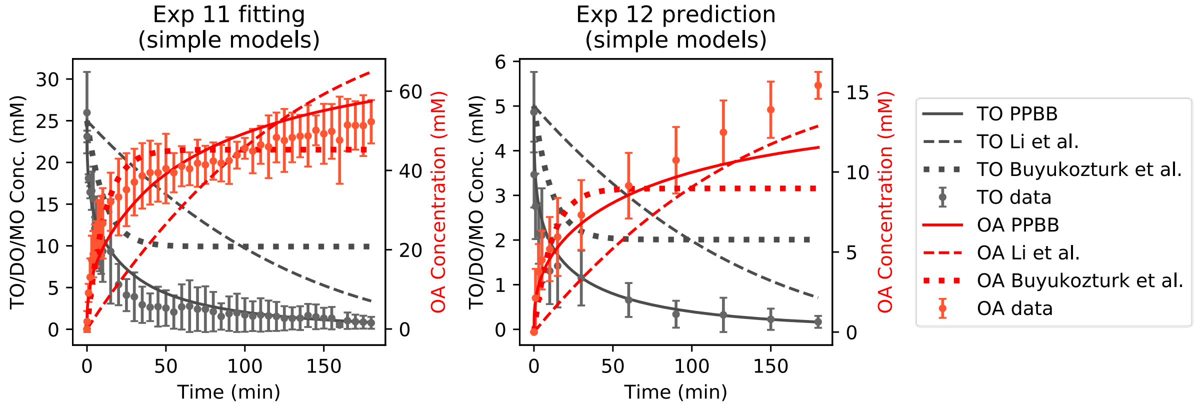


*Figure S7: Left - Fitting of lipolysis data from experiment 11 to models by Li et al. and Buyukozturk et al.; Right – Comparison of model predictions among the three models.*

*Table S8: Digestion coefficients from the Li et al. and. Buyukozturk et al. models fit to experimental data obtained at different micelle and enzyme concentrations. Units for k are mol $m^{-2}$ $min^{-1}$. Units for $k_{inh}$ are $min^{-1}$.*

| **Micelle/Enzyme** | ***k* (Li & McClements)** | ***k* (Buyukozturk et al.)** | **$k_{inh}$ (Buyukozturk et al.)** |
|---|---|---|---|
| Micelle: 3mM NaTDC:1mM EYPC Enzyme: 600U/mL | $8.77 \times 10^{-6}$ | $4.88 \times 10^{-5}$ | $3.99 \times 10^{-2}$ |

| | | | |
|---|---|---|---|
| Micelle: 12mM NaTDC:4mM EYPC<br>Enzyme: 600U/mL | $7.51 \times 10^{-6}$ | $2.09 \times 10^{-5}$ | $1.73 \times 10^{-2}$ |
| Micelle: 12mM NaTDC:4mM EYPC<br>Enzyme: 1800U/mL | $7.50 \times 10^{-6}$ | $4.94 \times 10^{-5}$ | $4.78 \times 10^{-2}$ |

### 8. Evaluation of different lipolysis mechanisms to improve monoolein profile prediction

As the MO profile was the least predictive by the proposed kinetic model, various attempts at improving the predictive ability of the model were made. For example, as noted previously, MO is produced at the oil-water interface upon hydrolysis of DO into 2-MO. However, 2-MO is not a substrate for pancreatic lipase, but it is for other lipases present in pancreatic extract (e.g., bile salt stimulating lipase) (Andersson et al., 1996). Concurrent to hydrolysis by these lipases, 2-MO also undergoes isomerization into 1-MO, which in turn is a substrate for pancreatic lipase. An attempt to capture this mechanism was made during the development of this model, but it yielded significantly more complex equations and four additional coefficients (not shown). The increased risk of data overfitting precluded obtaining a clear answer as to whether modeling the more complex MO mechanism could capture the kinetics of MO and DO profiles more accurately.

Other attempts to improve the MO profile fitting included considering a Langmuir adsorption model for partitioning of MO between the oil-water interface and the bulk solution. This change to the model resulted in worse fits as reflected in the SSND values generated (not shown). Another attempted change to the model was to consider not only digestion of MO at the interface but also that of MO in micelles. This digestion mechanism was assumed to follow the kinetics of an elementary reaction, i.e., to be proportional to the product of MO concentration in micelles ($C_{M_{aq}}$) and free enzyme concentration in solution ($C_E$). The addition of one extra coefficient again did not improve the fitting results (not shown). While the digestion mechanism of the aqueous MO is more likely to follow the saturable Michaelis-Menten kinetic model given that it is an enzymatic process, it was deemed unnecessary to incorporate considering that no appreciable improvement was achieved by the consideration of micellar MO being a substrate for digestion.

It is possible that discrepancies in MO profiles and model predictions could be explained by coalescence of particles during digestion. Several studies report coalescence of oil droplets following lipase addition and initiation of digestion (Day et al., 2014; Giang et al., 2015; Mun et al., 2007). However, coalescence during lipolysis was not considered in the model due to its stochastic nature and challenges associated with mathematically describing this complex process. In the interest of understanding the significance of coalescence on the lipolysis profile, we conducted experiments with conditions from experiment 2 where we monitored the evolution of droplets over time. We observed coalescence of particles in the first 10 min from the start of lipolysis followed by a decay in particle size over time (Figure S8).

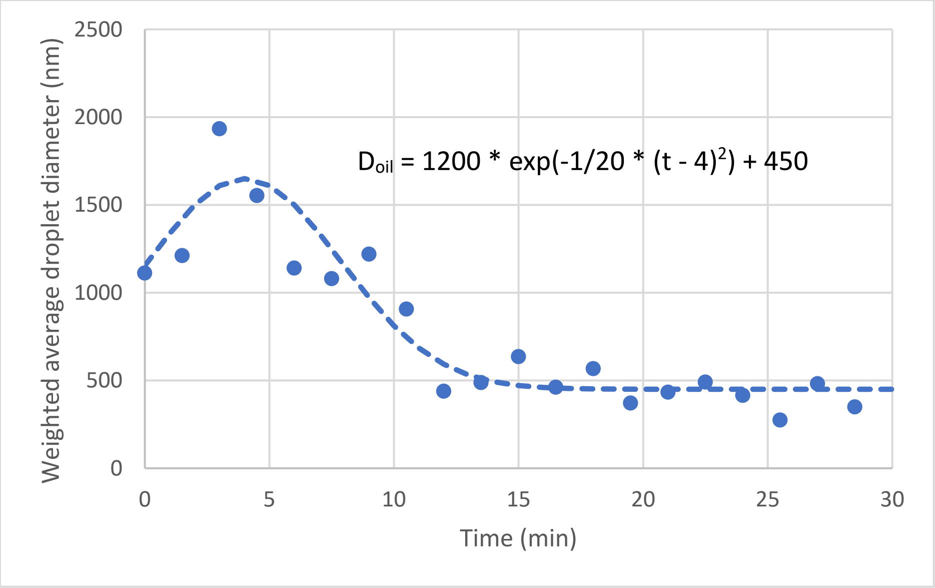


*Figure S8: Evolution of oil droplets during lipolysis of 25mM TO following conditions in experiment 2. $D_{oil}$ represents the emulsion diameter in nm and t represents the time since the start of digestion.*

We fit an exponential function to the data and used it to compute a variable number of oil droplets and consequently interfacial area, which were incorporated into the model. Model predictions considering vs. not considering coalescence are shown in Figure S9 relative to experimental results.

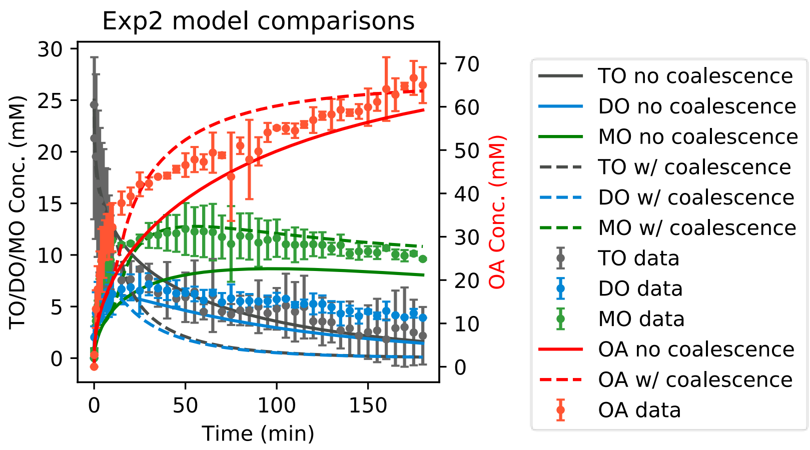


*Figure S9: Prediction of lipolysis profile for experiment 2 when discounting vs. accounting for droplet coalescence*

The results show an improved prediction with respect to MO when considering the coalescence of oil droplets during lipolysis and suggest that coalescence has a significant impact in lipolysis kinetics. However, incorporating the impact of coalescence resulted in overprediction of the rates of digestion of TO and DO. This is likely because the kinetic coefficients used were obtained by ignoring coalescence and are thus too large when accounting for coalescence. Because coalescence is stochastic and largely dependent on environmental conditions including solution volumes, mixing speeds, shear stress etc., it is difficult to model, in particular in a fashion that would likely translate to the *in vivo* environment.

### 9. Supplementary References

Andersson, L., Carriére, F., Lowe, M. E., Nilsson, A., & Verger, R. (1996). Pancreatic lipase-related protein 2 but not classical pancreatic lipase hydrolyzes galactolipids. *Biochim Biophys Acta*, *1302*(3), 236-240. https://doi.org/10.1016/0005-2760(96)00068-9

Brockman, H. L. (2000). Kinetic behavior of the pancreatic lipase-colipase-lipid system. *Biochimie*, *82*(11), 987-995. https://doi.org/10.1016/s0300-9084(00)01185-8

Capolino, P., Guérin, C., Paume, J., Giallo, J., Ballester, J.-M., Cavalier, J.-F., & Carrière, F. (2011). In Vitro Gastrointestinal Lipolysis: Replacement of Human Digestive Lipases by a Combination of Rabbit Gastric and Porcine Pancreatic Extracts [journal article]. *Food Dig*, *2*(1), 43-51. https://doi.org/10.1007/s13228-011-0014-5

Christiansen, A., Backensfeld, T., Denner, K., & Weitschies, W. (2011). Effects of non-ionic surfactants on cytochrome P450-mediated metabolism in vitro. *Eur J Pharm Biopharm*, *78*(1), 166-172. https://doi.org/10.1016/j.ejpb.2010.12.033

Couallier, E., Riaublanc, A., David Briand, E., & Rousseau, B. (2018). Molecular simulation of the water-triolein-oleic acid mixture: Local structure and thermodynamic properties. *J Chem Phys*, *148*(18), 184702. https://doi.org/10.1063/1.5021753

Day, L., Golding, M., Xu, M., Keogh, J., Clifton, P., & Wooster, T. J. (2014). Tailoring the digestion of structured emulsions using mixed monoglyceride–caseinate interfaces. *Food Hydrocolloids*, *36*(0), 151-161. https://doi.org/http://dx.doi.org/10.1016/j.foodhyd.2013.09.019

DebMandal, M., & Mandal, S. (2011). Coconut (Cocos nucifera L.: Arecaceae): In health promotion and disease prevention. *Asian Pacific Journal of Tropical Medicine*, *4*(3), 241-247. https://doi.org/https://doi.org/10.1016/S1995-7645(11)60078-3

Feeney, O. M., Williams, H. D., Pouton, C. W., & Porter, C. J. (2014). 'Stealth' lipid-based formulations: poly(ethylene glycol)-mediated digestion inhibition improves oral bioavailability of a model poorly water soluble drug. *J Control Release*, *192*, 219-227. https://doi.org/10.1016/j.jconrel.2014.07.037

Fernandez, S., Rodier, J.-D., Ritter, N., Mahler, B., Demarne, F., Carrière, F., & Jannin, V. (2008). Lipolysis of the semi-solid self-emulsifying excipient Gelucire® 44/14 by digestive lipases. *Biochimica et Biophysica Acta (BBA) - Molecular and Cell Biology of Lipids*, *1781*(8), 367-375. https://doi.org/http://dx.doi.org/10.1016/j.bbalip.2008.05.006

Giang, T. M., Le Feunteun, S., Gaucel, S., Brestaz, P., Anton, M., Meynier, A., & Trelea, I. C. (2015). Dynamic modeling highlights the major impact of droplet coalescence on the in vitro digestion kinetics of a whey protein stabilized submicron emulsion. *Food Hydrocolloids*, *43*, 66-72. https://doi.org/http://dx.doi.org/10.1016/j.foodhyd.2014.04.037

Hamilton, J. A., & Small, D. M. (1981). Solubilization and localization of triolein in phosphatidylcholine bilayers: a 13C NMR study. *Proc Natl Acad Sci U S A*, *78*(11), 6878-6882. https://doi.org/10.1073/pnas.78.11.6878

Koehl, N. J., Holm, R., Kuentz, M., Jannin, V., & Griffin, B. T. (2020). Exploring the Impact of Surfactant Type and Digestion: Highly Digestible Surfactants Improve Oral Bioavailability of Nilotinib. *Mol Pharm*, *17*(9), 3202-3213. https://doi.org/10.1021/acs.molpharmaceut.0c00305

Lewinska, A., Zebrowski, J., Duda, M., Gorka, A., & Wnuk, M. (2015). Fatty Acid Profile and Biological Activities of Linseed and Rapeseed Oils. *Molecules*, *20*(12), 22872-22880. https://doi.org/10.3390/molecules201219887

Minekus, M., Alminger, M., Alvito, P., Ballance, S., Bohn, T., Bourlieu, C., . . . Brodkorb, A. (2014). A standardised static in vitro digestion method suitable for food - an international consensus. *Food Funct*, *5*(6), 1113-1124. https://doi.org/10.1039/c3fo60702j

Montoya, C., Cochard, B., Flori, A., Cros, D., Lopes, R., Cuellar, T., . . . Billotte, N. (2014). Genetic architecture of palm oil fatty acid composition in cultivated oil palm (Elaeis guineensis Jacq.) compared to its wild relative E. oleifera (H.B.K) Cortés. *PLoS One*, *9*(5), e95412. https://doi.org/10.1371/journal.pone.0095412

Mun, S., Decker, E. A., & McClements, D. J. (2007). Influence of emulsifier type on in vitro digestibility of lipid droplets by pancreatic lipase. *Food Research International*, *40*(6), 770-781. https://doi.org/https://doi.org/10.1016/j.foodres.2007.01.007

Reis, P. M., Raab, T. W., Chuat, J. Y., Leser, M. E., Miller, R., Watzke, H. J., & Holmberg, K. (2008). Influence of Surfactants on Lipase Fat Digestion in a Model Gastro-intestinal System. *Food Biophys*, *3*(4), 370-381. https://doi.org/10.1007/s11483-008-9091-6

Tan, Y., Zhang, Z., Muriel Mundo, J., & McClements, D. J. (2020). Factors impacting lipid digestion and nutraceutical bioaccessibility assessed by standardized gastrointestinal model (INFOGEST): Emulsifier type. *Food Res Int*, *137*, 109739. https://doi.org/10.1016/j.foodres.2020.109739

ThermoScientific. (2022). *Certificate of analysis - Virgin olive oil (Lot Number: A0428595)*. https://www.thermofisher.com/document-connect/document-connect.html?url=https://assets.thermofisher.com/TFS-Assets%2FCCG%2Fcertificate%2FCertificates-of-Analysis%2FGEEL_41654_A0428595.PDF

Zhu, X., Ye, A., Verrier, T., & Singh, H. (2013). Free fatty acid profiles of emulsified lipids during in vitro digestion with pancreatic lipase. *Food Chem*, *139*(1-4), 398-404. https://doi.org/10.1016/j.foodchem.2012.12.060
